# Improving Growth Predictions in Aquaculture through an Improved Bioenergetics Model Incorporating Feed Composition and Nutrient Digestibility for Largemouth Bass (*Micropterus salmoides*)

**DOI:** 10.64898/2026.02.18.706619

**Authors:** Chunlin Chen, Lihua Song, Guoping Lian, Daoliang Li, Michael Short, Ran Zhao, Lianxiang Liu

## Abstract

Bioenergetics models serve as mechanistic tools to predict growth by linking energy intake, metabolic expenditure, and nutrient partitioning. However, traditional models rely primarily on gross energy (GE) intake, thereby oversimplifying the effects of feed composition and nutrient availability on fish growth. This work therefore proposed a refined bioenergetics model incorporating nutrient-specific digestibility coefficients (ADCs) and feed composition and tested using a compiled dataset (n=235; 165 for calibration and 70 for independent validation) and a field experimental dataset of largemouth bass (*Micropterus salmoides*). We first optimized parameters of a gross energy intake–based bioenergetics model, increasing R^2^ from 0.62 to 0.96 and thereby providing a calibrated foundation for subsequent refined model. The refined model demonstrated superior predictive performance on the compiled dataset (R^2^ = 0.97) with RMSE = 19.86 g and MAE = 10.31 g), representing reductions of 4.13% in RMSE and 19.98% in MAE and a 1.03% increase in R^2^ compared with the optimized GE-based model. In the field experiment, the refined model achieved high predictive accuracy (R^2^ = 0.98 and 0.97), whereas the optimized GE-based model showed poor performance (R^2^ = 0.33 and 0.06 respectively). This study is, to our knowledge, the first bioenergetics framework for largemouth bass that decomposes feed composition and nutrient-specific ADCs to compute macronutrient-resolved digestible energy, enabling formulation-aware growth prediction and nutrient-oriented optimization.

## 1. Introduction

Bioenergetics models outperform empirical growth curves, such as von Bertalanffy Growth Model (van Poorten & Walters, 2016), Logistic (Ma et al., 2018), and Gompertz (Sun & Wang, 2024), those are purely empirical fit that does not provide a causal or mechanistic link from diet, temperature and feeding to growth, and then Pütter model (Brunner et al., 2021), Thermal Growth Coefficient model (Jobling, 2003) add allometric or thermal structure but still lack explicit energy budgeting, whereas bioenergetics models encode ingestion, assimilation, maintenance, and excretion by partitioning dietary energy metabolic processes, enabling mechanistic interpretability and stronger prediction under diet, temperature, feeding changes (Chowdhury et al., 2013). Bioenergetics models have been widely used to support aquaculture management, including population dynamics estimation (Aljehani et al., 2023), fish growth tracking (Chahid et al., 2021), and feeding management (Karimanzira et al., 2016).

There have been several limitations of traditional bioenergetics models. They typically treat ingested energy as a homogeneous input, overlooking variations in macronutrient composition and digestibility across different feed formulations (Csargo et al., 2012; Hanson et al., 1997; Li et al., 2022; Portz & Cyrino, 2004). Parameters derived are mostly from low energy density feed (wild forage fish) that can no longer accurately reflect current industry practices (Li & Yakupitiyage, 2003; Rice et al., 1983; Rice & Cochran, 1984). Recent research demonstrated that the dietary macronutrient composition, specifically the levels and sources of protein, lipids, and carbohydrates influence not only growth efficiency but also key metabolic pathways, nutrient retention, and fish health (Portz & Cyrino, 2004), has brought growing attention to the effects of feed composition. Studies have further revealed that dietary macronutrient profiles and digestibility coefficients significantly affect nutrient retention and environmental impact, highlighting the limited biological resolution of energy-only models for precise dietary formulation (Dumas et al., 2010; Raposo et al., 2024). Protein is essential for somatic growth and tissue synthesis. Excessive intake or imbalanced energy supply may increase amino acid catabolism and nitrogenous waste, impairing both growth and water quality (Ekmann et al., 2013). Lipids, as dense energy sources, can spare protein for anabolic processes but may cause excessive fat deposition and metabolic disorders when overused, as observed in largemouth bass (Subhadra et al., 2006), gilthead seabream (Mongile et al., 2016), etc. Carbohydrates, though cost-effective, are variably utilised among species; high levels can reduce growth and nutrient retention in carnivorous fish such as largemouth bass (Amoah et al., 2008; Lin et al., 2018), and tambaqui (Sandre et al., 2017). Imbalanced macronutrient supply can also alter immune responses, hepatic metabolism, and enzyme activity related to energy and nutrient utilisation (Lin et al., 2018). These findings highlight the need to incorporate nutrient-specific responses into fish growth models to improve feed formulation development, support animal welfare, reduce costs of feed and water treatment, and minimise environmental impacts to support systems approaches for optimizing recirculating aquaculture systems (Zhang et al., 2023).

Previous studies reported different energy density of macronutrients (e.g., protein 23.6 kJ·g^−1^, lipid 39.5 kJ·g^−1^ and carbohydrate 17.2 kJ·g^−1^ (Csargo et al., 2012)) and wide ingredient-dependent ranges of ADC (e.g., protein 81.5-97.3%, lipid 50.6-98.2% (Qiu et al., 2022a)), demonstrating the significance of ingredients digestibility. Recent studies have promoted mechanistic bioenergetics models that integrate nutrient fluxes and physiological mechanisms to enhance predictive accuracy and applicability. For example, traditional bioenergetic model was extended by explicitly including the Energy and Protein fluxes, which can estimate fish growth by simulating fish body composition (Nobre et al., 2019), Dynamic Energy Budget models energy acquisition and allocation in organisms by separating biomass into reserve, structure and the reproduction compartments, which is limited by its complexity and high parameterisation requirements (Pecquerie et al., 2009). Nonetheless, many such models remain constrained by limited data and dependence on species-specific calibration, which limits their scalability across systems. This study aims to: 1) compile a comprehensive feed-growth dataset for *Micropterus salmoides* under modern intensive aquaculture conditions with high energy density feed; 2) optimize species-specific parameters of bioenergetics model to improve its prediction accuracy; 3) propose a refined modelling framework that integrates feed composition and nutrient-specific apparent digestibility coefficients (ADCs), enabling a macronutrient-based calculation of digestible energy intake.

This is, to our knowledge, the first attempt to explicitly combine feed composition breakdown with nutrient-specific ADCs in a bioenergetics framework for largemouth bass, parameterised and validated using both the largest comprehensive literature-derived dataset (235 records) and newly collected experimental data. This refined model structure enables better representation of feed composition effects on growth and provides a solid foundation for evaluating feed efficiency, nutrient utilisation, and waste output, while the dual-data design ensures generalisability while maintaining strong practical applicability.

## 2. Materials and Methods

### 2.1 Data Collection and Processing

A total of 235 sets of fish growth data were compiled from published studies of largemouth bass under diverse feeding and environmental conditions. Each record corresponds to an independent experiment, extracted from peer-reviewed journal articles and theses. Inclusion criteria are: 1) studies reporting detailed records of growth performance, 2) studies reporting detailed feed compositions (feed formulation, protein, lipid, energy density >12 kJ·g^−1^), 3) experiments with controlled growth conditions and feed conversion efficiency measurements.

ADCs for protein and lipid were extracted or estimated based on ingredient type, using representative values from digestibility trials reported in the literature. All values were normalized to consistent units (e.g., g for fish weight, % dry matter basis for feed composition, %/day for growth rate), and missing or conflicting data were treated using conservative interpolation or excluded if reliability was insufficient. The unified dataset enables the construction of a refined bioenergetics model that explicitly incorporates nutrient composition and digestibility efficiency for improved prediction of fish growth and feed utilisation. Detailed variables are shown in **Table1 (All tables can be found in appendix A)** below.

Besides, as shown in **Fig.1 (All figures can be found in appendix B)**, field experiments were conducted to validate the model as well, conducted in a recirculating aquaculture system (RAS) located in Hebei Province, China, consisting of two identical cylindrical tanks (1.5 m in diameter, 0.7 m water depth). Each tank was randomly stocked with 169 juvenile largemouth bass, initial body weight 15.1 ± 2.8 g. The two dietary treatments were as follows: diet A contains 53% crude protein, 8% of crude lipid and 19.35 kJ·g^−1^ gross energy, while diet B contains 43% crude protein, 12% of crude lipid and 19.63 kJ·g^−1^ gross energy. Fish were fed twice daily at an equal feeding rate based on biomass. Water quality parameters, including temperature, dissolved oxygen (DO), and pH, were continuously monitored throughout the 50 days experimental period. Growth performance was manually measured at regular intervals.

### 2.2 Baseline Bioenergetics Model

The baseline bioenergetics model applied in this study follows the classical energy budgeting framework commonly used in fish growth modelling (Rice et al., 1983). The model calculates the net energy balance of the ingested feed into three major pathways: metabolism, waste, and growth.

Specifically, the energy budget for non-reproducing fish can be summarised in the equation (1), in energy units (kJ g^-1^ d^-1^).:

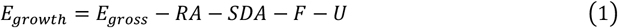

where *E*_*growth*_ is specific growth rate, *E*_*gross*_ is total energy from feed intake (kJ·d^-1^), *RA* is specific rate of standard metabolism and metabolism due to activity(kJ·d^-1^), *SDA* is the specific dynamic action rate, or energy required to digest food (kJ·d^-1^), *F* is specific egestion rate (kJ·d^-1^) and *U* is specific excretion rate (kJ·d^-1^).

The total energy of feed intake can be expressed as equation (2):

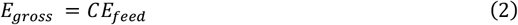

where *C* represents the amount of feed consumption(g·d^-1^), *E*_*feed*_ represents energy density of fish feed (kJ·g^-1^).

A previous study derived a respiration model for largemouth bass including standard metabolism and metabolic rate of activity(Rice, 1981), based on data from Beamish *et al*. (Beamish et al., 1970), as shown in equation (3).

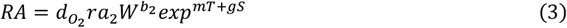

where 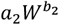 represents the maximum weight-specific standard respiration rate at the optimal temperature for respiration. Here, *a*_2_ represents the intercept for respiration and *b*_2_ represents weight dependence exponent for respiration. *e*^*mT*^, *e*^*gs*^ represent the temperature-dependent coefficient of respiration and the swimming speed-dependent coefficient of respiration, respectively. m and g are coefficients for temperature dependence of respiration and for swimming speed dependence of respiration respectively. Moreover, Rice used data from a variety of sources to construct a series of composite daily activity patterns and derived estimates of mean swimming speed for each composite pattern. This showed that little error is incurred by using a constant mean speed rather than variable speeds in modelling simulations(Rice, 1981). Units of respiration were converted from mg O_2_·g^-1^·h^-1^ to J·g^-1^·h^-1^ using an oxycalorific coefficient 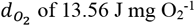 (Rice et al., 1983) and from J·g^-1^·h^-1^ to kJ·g^-1^·d^-1^ using a conversion constant *r*=0.024.

Specific dynamic action (SDA) is defined as the metabolic loss due to feed ingestion and digestion, and is assumed to be a constant proportion of the energy intake (Jobling, 1993), as shown in equation (4).

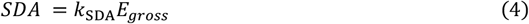

where *k*_SDA_ is a coefficient, which was cited as 14.2% of the energy consumed(Beamish, 1974; Rice et al., 1983).

Energy losses through egestion (F) and excretion (U) have also been estimated as constant proportions of the energy consumed, as expressed in equation (5) and (6), respectively.

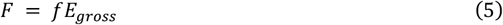

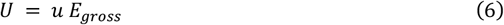

where f and u are coefficient of egestion (faecal loss) and excretion (non-faecal), respectively. For largemouth bass, Beamish estimated egestion to be 10.4% of total energy consumed, and estimated excretion as 7.9% of total energy consumed (Niimi & Beamish, 1974).

In summary, the state equation of baseline bioenergetics model can be expressed as equation (7).

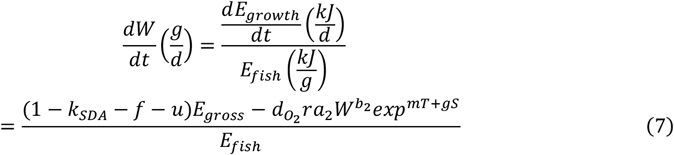

In this gross energy based model (set as baseline model), the values of the parameters are shown in **Table 2**.

The baseline model operates under the following assumptions: 1) The energy intake and utilisation are mainly determined by digestible protein and lipid, assuming carbohydrate contribution is minimal as largemouth bass is carnivorous species. 2) Total energy intake is partitioned into respiratory metabolism, egestion and excretion, growth, and SDA. 3) The energy density of the fish body is assumed to remain constant, at 4.184 kJ·g^−1^. These assumptions, while useful for simplicity and generalisation, limit the model’s ability to accurately capture nutritional effects and feed-specific performance outcomes under diverse feeding regimes

### 2.3 Refined Bioenergetics Model

While the baseline model’s calculation of gross energy intake (*E*_*gross*_) assumes homogeneity of energy sources, neglecting variation in digestibility among feed ingredients, the refined model was extended to a composition-aware intake (composition × ADCs) framework, incorporating dietary protein, lipid, and carbohydrate fractions, and associated ADCs of different ingredients. Egestion is no longer considered as digestible energy is calculated directly, which decrease error caused by estimating egestion. **Fig.2** illustrates the difference between the baseline model and the proposed refined model, highlighting the addition of feed composition and ADCs prior to energy allocation. Consequently, the overall refined bioenergetics model is expressed in the equation (8).

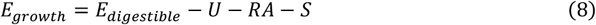

where *E*_*fish*_ refers to the energy density of the fish body, usually set as a constant of 4.184 kJ·g^-1^ (Rice et al., 1983).

The ADC of different energy sources can be calculated by equation (9).

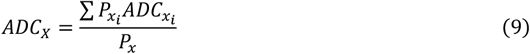

where *x* ∈ *X* represents crude protein (*p*), crude lipid (*l*) and carbohydrate (*CHO*), *P*_*x*_ represents the total content of 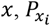 represents the content of component *i* from different sources, *ADC*_*X*_ represents ADC of component *x*, which is denoted as *D*_*p*_, *D*_*l*_ *and D*_*CHO*_, and *i* ∈ *I* represents ingredients of component x. Specifically, digestible energy (*E*_*digestible*_) is derived from the digestible portions of protein, lipid, and carbohydrate based on their respective ADCs, as shown in equation (10). By decoupling nutrient-specific digestibility and energy input, the model can capture differences between feeds with equal protein or lipid content but differing quality or ingredient source.

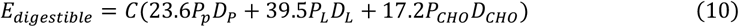

where *C* represents feed consumption, *P*_*p*_, *P*_*l*_, *P*_*CHO*_ (%) represent proportion of crude protein, lipid and carbohydrate in the fish feed, and *D*_*P*_, *D*_*L*_, *D*_*CHO*_ represent ADCs of those macronutrients. 23.6, 39.5 and 17.2 (kJ·g^−1^) are the combustion values of protein, lipid and carbohydrate (Csargo et al., 2012; Nobre et al., 2019).

In the refined model, the non-faecal energy loss (*U*), representing energy lost through urinary and gill excretion, is tied to the digestible portion of feed energy as it arises from metabolized and absorbed nutrients, expressed as equation (11).

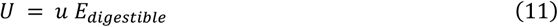

where *u* is the coefficient for excretion.

Notably, as multiple studies have consistently shown that largemouth bass predominantly utilise amino acids rather than glucose or fatty acids as energy substrates, and given that, as a carnivorous species, they exhibit high protein utilisation efficiency but low carbohydrate digestibility. This model therefore considered only protein and lipid as energy sources, excluding carbohydrates (X. Li et al., 2020; Li & Wu, 2019). Moreover, although fish energy density is difficult to measure frequently, but previous study provides valuable empirical estimation method for growing fish (Canale & Breck, 2013), as shown in equation (12). In summary, the state equation of the refined bioenergetics model can be expressed as equation (13).

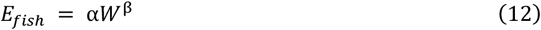

Coefficients α, β are empirical constants. It was reported that α, β are 5.01 and 0.046 respectively [20]. Overall, the refined bioenergetics model is shown as follows:

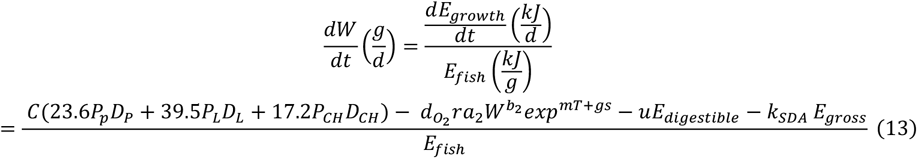

The refined model explicitly accounting for nutrient composition and digestibility, enabling more precise and feed-specific growth predictions. In this model, it assumes ADC values extracted from literature are assumed to be representative for similar ingredient categories across studies.

### 2.4 Model Calibration and Evaluation Strategy

Following the theoretical framework, this section details the model calibration. Key parameters for energy partitioning and metabolic loss were estimated using a comprehensive dataset of 235 experimental records to ensure biological accuracy and practical applicability. The model is a generic bioenergetics framework designed for applicability across fish species. To confirm its effectiveness, it was specifically applied to and validated through a case study on largemouth bass, assessing its predictive accuracy and biological relevance under real-world aquaculture conditions.

#### 2.4.1 Parameter Optimization

The model parameterisation for a given species is also important for the consistent and general model application. Initially, as shown in Table 2, the baseline model was implemented using initial parameter values cited in classical bioenergetics literature [13]. These were primarily derived from studies conducted on largemouth bass fed with low energy density diets, such as lake emerald shiners (*Notropis atherinoides*)-based diets (Rice et al., 1983). To improve the model accuracy and adapt to current feeding practices, this study re-estimated key model parameters using the compiled dataset with 235 records. This study employed a constrained nonlinear least-squares optimization using fmincon in MATLAB, integrated with a multi-start global search strategy. Initial parameter sets were generated via Latin Hypercube Sampling (LHS) within physiologically plausible bounds to ensure global convergence. Six physiological parameters *a*_2_, *b*_2_, *m, k* _*SDA*_, *f* and *u*, were estimated by minimising the sum of squared errors (SSE) between predicted and observed body weights. The bounds of parameters *a*_2_, *b*_2_, *m, k*_*SDA*_, *f* and *u*, based on some empirical data, are [0,1], [-0.6,-0.1] (Beamish et al., 1970), [0.02, 0.09] (Rice et al., 1983), [0.08,0.2] (Beamish, 1974; Rice et al., 1983; Tandler & Beamish, 1981), [0.08 0.2], [0.05 0.15] (Whitledge & Hayward, 1997), ensuring the optimization results were physiologically realistic and overfitting avoided. To prevent overfitting and promote parameter regularity, an L2-regularisation term was added, proportional to the sum of the squared parameter values, scaled by a regularisation coefficient *λ*. This framework yielded robust and interpretable parameter estimates across multiple initialisations. The loss function is expressed in equation (14).

The loss function is as follows:

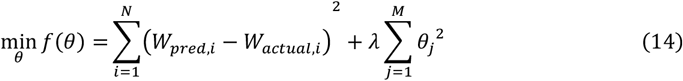

where *W*_*pred,i*_ and *W*_*actual,i*_ represent the predicted and observed body weight for sample *i, θ*_*j*_ denotes the *j*-th model parameter to be optimized, λ is the regularisation coefficient controlling the penalty strength on parameter values, N is the number of samples and M is the number of parameters.

#### 2.4.2 Sensitivity Analysis

Sensitivity analysis was conducted using two complementary approaches to evaluate the influence of model parameters on output variables. First, a local One-At-a-Time (OAT) method was applied, in which each parameter was varied individually while holding others constant, to assess its direct effect on model outputs. Subsequently, a global sensitivity analysis was performed based on Monte Carlo sampling combined with Pearson correlation analysis. Many parameter sets were generated through random sampling, and the model was executed for each set. Pearson correlation coefficients between each input parameter and the model output were calculated, then squared and normalized to indicate the relative importance of each parameter. This two-tiered strategy enabled both the identification of individual influential parameters and the detection of potential interactions and non-linear effects across the model.

#### 2.4.3 Model Validation and Evaluation

The compiled dataset of 235 samples was randomly divided into a training subset and a test subset at a 7:3 ratio. The training subset (n = 165) was used for parameter optimization, while the independent test subset (n = 70) was used in both baseline and refined bioenergetics model to validate model performance. Field experimental data were employed for further comparative validation of the model, providing a robust test platform for assessing model generalisability.

Specifically, model performance was assessed by comparing predicted body weight to the actual weight. Here, two derived performance metrics, specific growth rate (SGR) and feed conversion ratio (FCR), were included as secondary evaluation metrics, as expressed in equation (15) and (16), respectively.

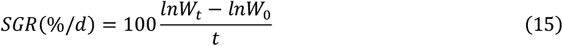

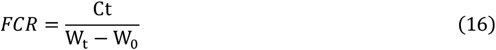

where *W*_*t*_, *W*_0_ represent the body weight (g) at time *t* and at the initial time (day 0), respectively; *C* represents feed consumption (g·g^-1^·d^-1^), *t* is duration of feeding period in days.

Model performance was assessed using metrics of root mean square error (RMSE), mean absolute error (MAE), and coefficient of determination (R^2^). Additionally, residual plots were examined to identify potential systematic biases or heteroscedasticity.

## 3. Results

### 3.1 Baseline Model Simulation

The baseline bioenergetics model was initially implemented using literature-derived parameters(Rice et al., 1983), primarily estimated from early studies using fish-based diets. However, when applied to the compiled dataset featuring high-energy feeds, it showed poor predictive accuracy with RMSE = 60.91 g, MAE =41.51 g, R^2^ = 0.62, showing significant overestimation on fish weight. as shown in **Fig.3(a)**. The derived metrics SGR and FCR were also poorly estimated, with R^2^= - 0.89 and R^2^= - 3.29. SGR is generally overestimated, while FCR is underestimated, as shown in **Fig.3(b-c)**. Additionally, systematic biases were observed, as shown in **Fig.3(e-f)**, the mean residual of W, SGR, and FCR are 41.51, 0.84 and -0.51, respectively. The results indicate that the cited parameters failed to accurately capture the fish growth dynamics observed in the compiled dataset.

### 3.2 Parameter Optimization and Sensitivity Analysis of Baseline Bioenergetics Model

#### 3.2.1 Sensitivity Analysis of cited parameters

Sensitivity analysis using both local OAT (**Fig.4 (a)**) and global Monte Carlo (**Fig4 (b)**) sampling approaches consistently identified the allometric coefficient for metabolic rate (*b*_2_), as the most influential parameters affecting the predicted weight significantly. Parameters *a*_2_ and *m* showed moderate sensitivity, while *k*_*SDA*_, *f* and *u* had limited impact, and the parameter *g* showed negligible effect across both methods. The high sensitivity of *b*_2_, *a*_2_, and m indicate their importance for calibration. In contrast, parameters such as g can be fixed during optimization to reduce model dimensionality without reducing accuracy. Overall, calibration efforts should prioritise *b*_2_, followed by *a*_2_, *m, k*_*SDA*_, *f* and *u*.

#### 3.2.2 Parameter Optimization

Prioritised parameters of the foregoing section were best simulated using a multi-start global optimization with fmincon. The objective function converged stably, satisfying the first-order optimality condition. The optimized model parameters are shown in **Table 3**.

As can be seen in **Table 4**, the results show that the optimized parameters significantly improved the prediction accuracy in compiled dataset, the RMSE and MAE decreased to 20.68 g and 12.37 g, respectively. The R^2^ increased to 0.96, and the residuals of SGR and FCR were decreased by 90.80% and 83.64%, respectively. **Fig.3** further illustrates improvements in predictive accuracy (a-c) and reductions in systematic biases (e-f). These results demonstrated the optimized parameters of baseline bioenergetics model more accurately captured the growth dynamics of largemouth bass under high-energy-density diets in the compiled dataset.

As shown in **Fig.5**, applying the baseline bioenergetics model with optimized parameters to the experimental dataset yielded significant improvements in prediction accuracy for both diets, compared to the results obtained using cited parameters, RMSE decreased from over 14 g to lower than 4.5 g. However, the R^2^ values remained low (near zero), indicating that despite to improve simulation capacity, the model still fails to adequately capture key features of the observed growth. This suggests that further refinement of the model structure or input parameters may be required.

#### 3.2.3 Sensitivity analysis of optimized parameters

As illustrated in **Fig.6**, both local and global sensitivity analyses consistently identify *b*_2_ as the most influential parameter after optimization, and *g* exhibited minimal sensitivity. In contrast, it also displayed the redistribution of parameter contributions, with increased relative importance for parameters *a*_2_, *k*_*SDA*_, *f, m* and *u*, implying a more balanced parameter influence structure. The model with optimized parameters demonstrates refined robustness while reducing overreliance on a single dominant parameter and retaining sensitivity to key biological processes.

### 3.3 Refined Bioenergetics Model

The baseline bioenergetics model exhibited systematic overestimation in body weight predictions and generated notable fitting errors for SGR and FCR. These discrepancies suggest that the model, originally developed using fish-based diets, fails to capture the metabolic dynamics associated with high-energy density commercial feeds represented in the compiled dataset. Following the parameter optimization, the baseline model remains unable to capture the influence of feed nutrient composition on fish growth.

To tackle these limitations, a refined model is introduced in the following section to account for differences in energy contribution from various dietary components and their respective proportions and digestibility.

#### 3.3.1 Nutrient Digestibility Profiles of Feed Ingredients

To improve the estimation of metabolically available energy, the refined model incorporated ingredients composition and apparent digestibility coefficients derived from literature and feeding trials. Table 1 lists a comprehensive range of proximate nutrient contents and ingredient inclusion levels derived from published literature and experimental records. Key nutritional parameters encompass crude protein (37.20-56.00%), crude lipid (3.30-23.30%), and gross energy (12.51-22.10 kJ·g^−1^). Ingredient-level data cover both animal- and plant-based sources, including fish meal (0.00-55.60%), soybean meal (0.00-55.10%), poultry byproduct meal (0.00-52.50%), and others. **Table 5** summarises the representative ADCs applied to key feed components within the dataset. These coefficients were used to compute digestible energy contributions from protein, lipid, and carbohydrate fractions, thereby allowing a more realistic simulation of energy intake and partitioning under high-energy diet conditions.

#### 3.3.2 Adjustment of Fish Energy Density and Swimming Speed

As shown in **Fig.7**, four approaches for estimating fish energy density were compared. Method (a), which requires detailed body composition data, was excluded due to practical limitations. The baseline model used a fixed value of 4.184 kJ·g^−1^, while the refined model adopted a power function after comparison with a piecewise alternative. The power function model provided the best performance, with R^2^ increasing from 0.90 (constant), to 0.95 (piecewise), and 0.96 (power function). Correspondingly, RMSE, MAE, and residuals for SGR and FCR were all reduced, indicating refined predictive accuracy.

After the model refinement, parameter sensitivity analysis was also conducted. **Fig.8** illustrates that parameter *b*_2_remained the most influential parameter on fish weight prediction in both the OAT and global sensitivity analyses, indicating their key roles in metabolic modelling. Compared to the original framework, the ranking of parameter sensitivity remained largely consistent, suggesting that the refined model retained the core sensitivity characteristics while incorporating refined structural mechanisms.

#### 3.3.3 Validation with Compiled Dataset

As summarized in **Fig.9** and **Table 6**, the refined model shows high predictive accuracy (R^2^=0.97) on the compiled dataset. As compared to the baseline model with optimized parameters, the refined model shows a slight improvement in weight prediction, the RMSE and MAE slightly decreased by 4.13% and 19.98%, respectively, while the R^2^ remained consistently high (0.96 to 0.97), indicating the model maintained strong predictive capability. However, the mean residuals for SGR and FCR shifted from 0.08 to -0.20 and from -0.09 to 0.17, respectively, highlighting the limitations of the refined model in accurately capturing the growth rate and feed conversion dynamics within the compiled dataset. Overall, the refined bioenergetics model, by incorporating literature-based feed formulations and ADCs-based energy intake, provides reliable predictions of fish growth and effectively characterises digestible energy utilisation pathways.

#### 3.3.4 Validation with Field Study

Moreover, field experiments with two diets were carried out to validate the effectiveness of the refined model. As shown in **Fig.10** and summarised in **Table 7**, as compared to the baseline bioenergetics model, the refined model significantly improves the predictive accuracy, as evidenced by reductions of over 80% in both RMSE and MAE, and R^2^ increases from 0.06 to 0.97 (CP 43%), and 0.33 to 0.98 (CP 53%). These results indicate that the refined model not only minimised prediction errors but also captured the growth dynamics of largemouth bass with high fidelity across both nutritional treatments. Consequently, the refined structure enhances the model’s applicability to contemporary aquaculture diets and improves its predictive fidelity for both growth and nutritional efficiency.

## 4. Discussion

Based on a baseline bioenergetics model, this study focuses on calibrating model parameters and updating the model framework. The parameter calibration successfully captured the growth dynamics of largemouth bass under high energy density commercial diets, agreeing with the physiological traits observed under intensive aquaculture conditions. Following that, the model was refined by incorporating empirical data of ingredients, feed formulations, and ADCs. These enhancements, informed by empirical and literature-based data, contributed to a more comprehensive framework capable of simulating the effects of feed formulation on growth outcomes. The refined model accurately reproduced growth trajectories from both published and field-collected datasets, underscoring its predictive reliability. Overall, the results demonstrate that the revised model, with its integrated consideration of feed digestibility and nutrient composition, offers a robust tool for predicting growth responses in largemouth bass. Furthermore, it holds considerable promise for future applications in precision feed formulation, enabling more efficient and sustainable aquaculture practices.

### 4.1 Parameter Sensitivity and Calibration

Previous studies recognized that respiration-related parameters, *b*_2_, *a*_2_ and *m*, are dominant drivers of model variability [13,47]. These parameters fundamentally regulate metabolic expenditure by scaling respiration with body mass and temperature, thereby exerting strong impact on growth predictions (Walters & Essington, 2010). However, after calibration using the compiled dataset collected under controlled aquaculture conditions, the relative sensitivity of respiration-related parameters declined, while postprandial and excretory costs are increasingly captured by coefficients related to SDA (*k*_*SDA*_), egestion (*f*) and excretion (*u*) rather than being implicitly absorbed into respiration. Biologically, this re-ranking reflects a transition in the primary constraints on growth: under relatively thermally stable farming conditions (25-29 ℃), respiration costs are reduced (Díaz et al., 2007). The reduction of *a*_2_ related to basal metabolic loss and increase of allometry *b*_2_ suggests a decreasing dependence on body mass (Bajer et al., 2004; Beamish et al., 1970). Empirical measurements support this outcome: *Q*_10_ = *e*^10*m*^ values for largemouth bass range from 1.37-2.00 within 26-32 °C, with optimum growth temperatures around 27.5 °C (Rice et al., 1983). The recalibrated parameter *m* yielded a *Q*_10_ =1.32, indicating reduced thermal sensitivity, an expected pattern in controlled systems with slight temperature fluctuations (Walters & Essington, 2010).

Reported SDA values for largemouth bass range from 10.5% on a dry pelleted diet containing 35.8% protein (Tandler & Beamish, 1979), 11.3% on a similar diet with 4.72 kcal·g^−1^ gross energy (Tandler & Beamish, 1981), to 14.19% (Beamish, 1974) and 16.3% (Tetzlaff et al., 2010) for a diet of freshly thawed lake emerald shiners, but SDA typically does not exceed 25% of ingested energy (Tandler & Beamish, 1979). In this study, the calibrated *k*_*SDA*_ = 0.20, which lies within the biologically plausible upper bound yet is higher than most previous estimates. We attribute this elevation primarily to dietary composition and energy density: compared to fish-based diets (approximately 15%-20% protein, 2%-20% lipid, 3 to 6 kJ·g^−1^, commonly ≈ 4.184 kJ·g^−1^, wet-weight basis), protein-dense diets (>40% protein and >12 kJ·g^−1^) increase the postprandial biochemical costs of protein digestion and assimilation, such as transamination, deamination, protein synthesis and turnover, thereby elevating post-feeding oxygen consumption and therefore SDA (Bucking, 2017; Guo et al., 2012; Secor, 2009). Consistent with this mechanism, experiments in largemouth bass show that SDA scales positively with dietary protein level and meal energy. For a given energy intake, SDA is highest on 100% protein and lowest on 100% carbohydrate diets, and increases with ration size and body mass (Tandler & Beamish, 1980). Convergent evidence across fishes further indicates that raising dietary protein increases oxygen demand, SDA magnitude and, in some cases, the SDA coefficient, consistent with protein’s higher thermic effect (Guo et al., 2012; Secor, 2009).

Protein-dense diets also increase excretory nitrogen loss, as showed in the quantitative relationship between nitrogen excretion (Y) and nitrogen intake (X), which can be described as Y=A (basal metabolism)+BX (Savitz et al., 2010). Together with increased nitrogenous excretion and associated ion-transport or acid-base regulation costs (Savitz et al., 2010; Tidwell et al., 1996). The energy equivalents for ammonia-nitrogen and urea-nitrogen are 5.94 calˑmg^−1^ and 5.51ˑcal mg^−1^, respectively (Elliott, 1976). Intake-side identifiability improved after integrating composition-specific ADCs, reducing compensatory effects previously absorbed by respiration coefficients.

Together, these results indicate that while respiration-related parameters remain the most influential, their relative dominance decreases in controlled, nutrient-rich conditions, with SDA and nitrogen-related excretory losses emerging as additional key sources of model variability. From a practical perspective, this highlights the importance of optimizing protein quality and digestibility to improve feed efficiency and reduce waste, complementing the traditional focus on maintaining suitable thermal environments. Nonetheless, the generality of these findings is uncertain. This higher SDA may also partly reflect limitations of the calibration dataset, which primarily included intensively fed fish under stable aquaculture conditions. In pond systems with greater thermal/feeding fluctuations or in populations with different metabolic backgrounds, respiration may regain higher sensitivity (Q10 approaching 2.0) (Díaz et al., 2007). Future work should therefore test whether the changed body-mass and temperature dependence observed here can be verified through respirometry trials, and whether parameter shifts reflect short-term acclimation or long-term genetic adaptation.

### 4.2 Refinement of Model Framework

Prior studies have advanced bioenergetics applications in production management (Csargo et al., 2012), feeding strategy (Li et al., 2022; Ranney et al., 2024), quantifying environmental effects (Niimi & Beamish, 1974), and species-specific parameter calibration (Fantini et al., 2021). Current research has shifted toward improving production efficiency through ingredient substitution (He, Li, et al., 2020), feed formulation optimization and ration management (Aizam et al., 2018; Li et al., 2022), yet these trail-based approaches remain labour-intensive and time-consuming. This motivates an update modelling framework that can capture feed-composition effects directly, reducing reliance on costly iterative trials. One promising direction is refining nutrient utilisation pathways. The refined model re-expressed gross energy intake as macronutrient-resolved digestible energy (DE), defined as the sum of digestible contributions from protein, lipid, and carbohydrate (with energy density of 23.6, 39.5, and 17.2 kJ g^−1^, respectively (Csargo et al., 2012)) and digestibility values compiled from over 20 studies on common ingredients (e.g., fish meal, soybean meal, wheat flour, and corn gluten) (Cui et al., 2022; Qiu et al., 2022b). Mechanistically, this matters because protein carries higher SDA costs and nitrogen excretion than lipid or carbohydrate; consequently, diet composition, not only total energy alone, regulates the partitioning of energy and growth. Consistent with this rationale, published studies report wide ingredient-dependent ranges of ADC (e.g., protein 81.5-97.3%, lipid 50.6-98.2%) (Portz & Cyrino, 2004; Qiu et al., 2022b; Tidwell et al., 2005), which baseline models implicitly ignore. This supports the robustness of the updated model structure. In addition, the proposed framework is supported by similar developments cross other fish species. For instance, an integrated energy-protein flux model extends traditional bioenergetics by explicitly tracking energy and protein fluxes with practical inputs (feed intake and composition, temperature, initial weight, and ADCs) to dynamically predict growth of gilthead seabream (Nobre et al., 2019). Evidence from Nile tilapia shows that there are direct relationship between digestible energy intake and energy gain, as well as between digestible protein intake and protein gain (Raposo et al., 2024). Other models have also introduced feed performance to compare diets with different compositions for *Sparus aurata*, supporting the inclusion of feed quality effects in growth predictions (Libralato & Solidoro, 2008). These cross-species results support the composition-resolved, digestible intake (composition × ADCs) approach for formulation optimization and scenario testing.

Furthermore, we also adjusted the common assumption of constant fish energy density. Energy density in largemouth bass changes with growth and body composition (reported ranges ≈ 4.4-5.5 kJ g^−1^ (Csargo et al., 2012),, and even 3.5-7.5 kJ g^−1^ (Glover et al., 2011)). Previous studies have modelled energy density as a function of body weight, including a two-phase linear model with a breakpoint at 174 g (Glover et al., 2011), and body-composition regressions using lipid, protein, and ash (Breck, 2011), and power functions (Canale & Breck, 2013). In this study, we adopted a mass-dependent power function for energy density, which better captures its ontogenetic dynamics and improved predictive accuracy. Nevertheless, energy density is also influenced by diet quality and environmental conditions (Breck, 2011; Johnson et al., 2017), which are not yet fully captured.

The refined model was validated using both a literature-based dataset and an independent field dataset, achieving R^2^ values of 0.95 and 0.97, respectively, indicating that a composition-aware intake (composition × ADCs) together with mass-dependent energy density captures diet effects across formulations. The proposed model, despite its refined structure and predictive accuracy (R^2^=0.96), maintains an acceptable level of data complexity requires initial body weight, temperature, feeding rate, ingredient-specific ADCs and feed composition, and remains applicable in production settings like nutrient regulation and feed ingredient substitution with limited inputs. FCR and SGR are essential indicators of feed efficiency and growth performance (Bright et al., 2005; Guo et al., 2019). Although derived from predicted body weight and feed intake, decreased prediction residuals reflects a more physiologically consistent representation of nutrient allocation and metabolic balance.

### 4.3 Applications, Limitations, and Improvement

The updated model enables scenario testing across dietary formulations due to the refined framework that integrates nutrient-specific ADCs. In practise, it also allows 1) estimation of ammonia excretion (TAN) from nutrient intake and nitrogen partitioning to support water quality management (Endut et al., 2014), and 2) feed formulation optimization, by evaluating the performance of different ingredient combinations and nutrient ratios under specific farming scenarios (Aizam et al., 2018). These uses target operational decisions, nutrient regulation, ingredient substitution, and cost–performance trade-offs, without requiring extensive trial-and-error.

Limitations remain. Accuracy in production settings can be constrained by proprietary formulations (incomplete ingredient disclosure) and situation-dependent conversion efficiencies (variation in digestibility and post-absorptive costs with size, temperature, processing, and ingredient source). These factors introduce parameter uncertainty that is not fully resolved by the current specification. To address this, it is suggested that developing an empirical, parameter-based efficiency mapping layer that infers effective nutrient conversion efficiencies from routinely available covariates, proximate composition (crude protein, lipid, carbohydrate), environmental variables, fish physiological traits, and digestibility metrics without requiring full ingredient disclosure (Cho, 1992). Such a layer would allow: 1) forward prediction of ADCs from basic feed profiles, and 2) potentially enable reverse optimization of feed composition based on target ADCs or performance outcomes. In turn, this would improve robustness for commercial deployment and shift the model from a generic energy-budgeting tool toward a nutritional decision platform that supports ingredient substitution, feed selection, and feeding-strategy refinement. Additionally, a scalable and integrative bioenergetics framework for commercial intensive aquaculture systems that bridges feed formulation, nutrient utilisation, and waste release control is required to enable real-world application (Karimanzira et al., 2016). Ensuring the feasibility of such models in industrial RAS settings will support economic viability while promoting environmentally sustainable aquaculture practices.

## 5. Conclusion

This study first compiled a literature-derived dataset of largemouth bass in intensive aquaculture with 235 records for model calibration and validation. The calibrated respiration parameters shifted to *a*_2_ =0.3393, *b* =00.3486, *m* =0.028, while *k*_*SDA*_ =0.20, and *u* =0.15 increased, which is interpreted as scenario-specific parameter reallocation consistent with protein-dense diets and thermally stable conditions. On the compiled literature dataset, parameter optimization for the baseline model reduced RMSE from 60.91 to 20.68 g (decreased by 66%) and MAE from 41.51 to 12.37 g (decreased by 70%), increasing R^2^ from 0.62 to 0.96. The study also presents the first bioenergetics framework for *Micropterus salmoides* that decomposes feed composition and nutrient-specific ADCs to refine macronutrient-resolved digestible energy. Results of validation show that the refined model achieved R^2^=0.97 on the literature-based dataset with RMSE = 19.86 g and MAE = 10.31 g (vs. the optimized-baseline model: RMSE decreased by 4.13%, MAE decreased by 19.98%, and R^2^ increased by 1.03%). On the independent field dataset with two isoenergetic but compositionally different diets, the refined model reached R^2^=0.98, 0.97 with RMSE = 0.76, 0.82 g and MAE = 0.66, 0.69 g respectively, whereas the baseline with optimized parameters performed poorly (R^2^< 0). Overall, refining intake as macronutrient-resolved DE and using mass-dependent energy density provide accurate, generalisable predictions across formulations, enabling formulation design, nutrient management (including TAN estimation), and scenario testing in intensive aquaculture.

## Data Availability Statement

The data that support the findings of this study are available from the corresponding author upon reasonable request.

## Acknowledgments

This work was supported by the UK-China Agritech Challenge: Aquaculture 4.0 -- Advancing Digital Precision Aquaculture in China (ADPAC) (**Project Reference:** BB/S020896/1) and a Research Scholarship from the China Scholarship Council.

## i. Appendix A: Tables

**Table 1.**
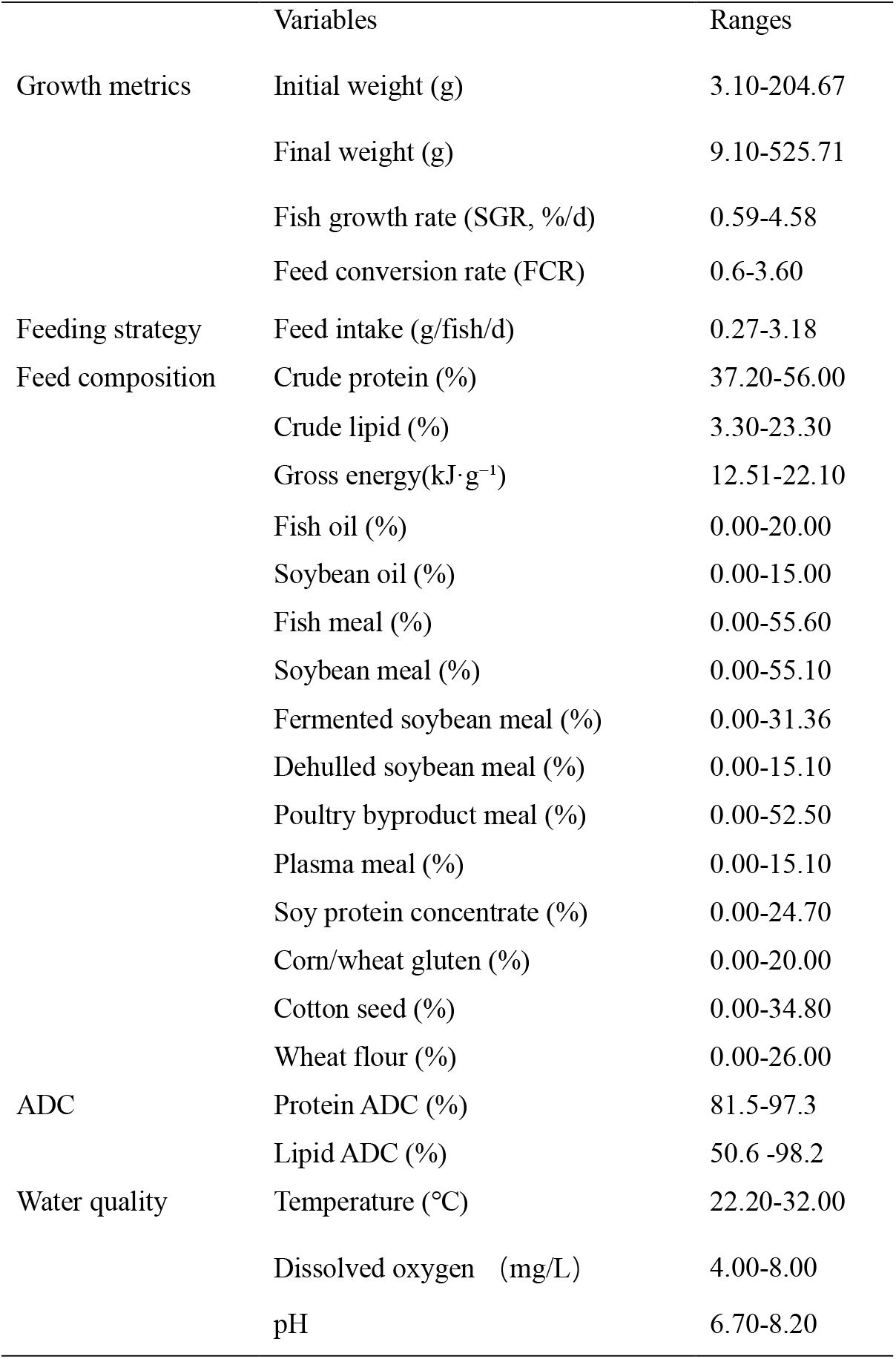
Collected variables from published literature for the model dataset.

**Table 2.**
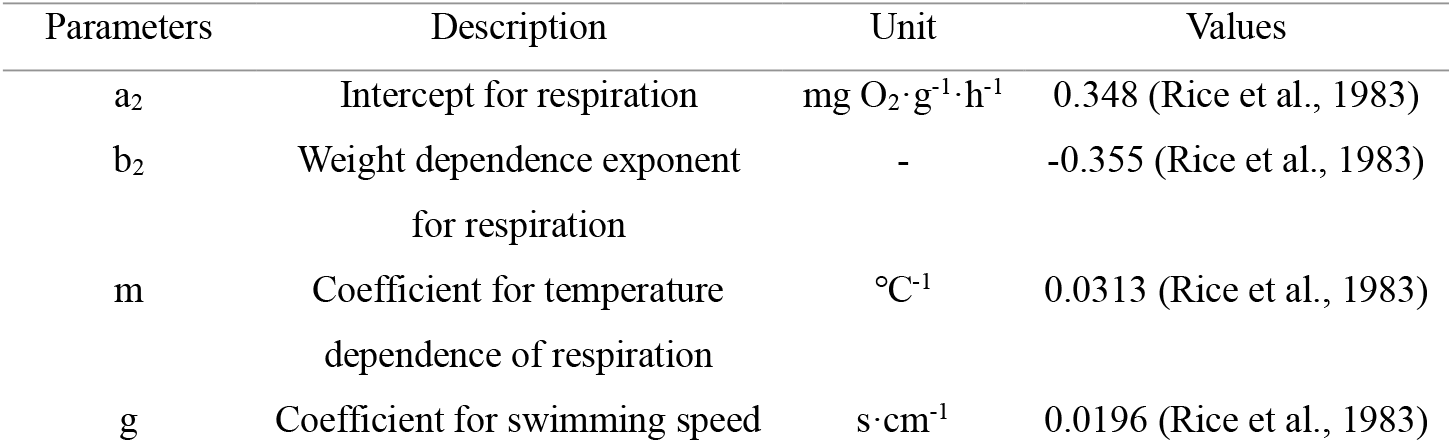
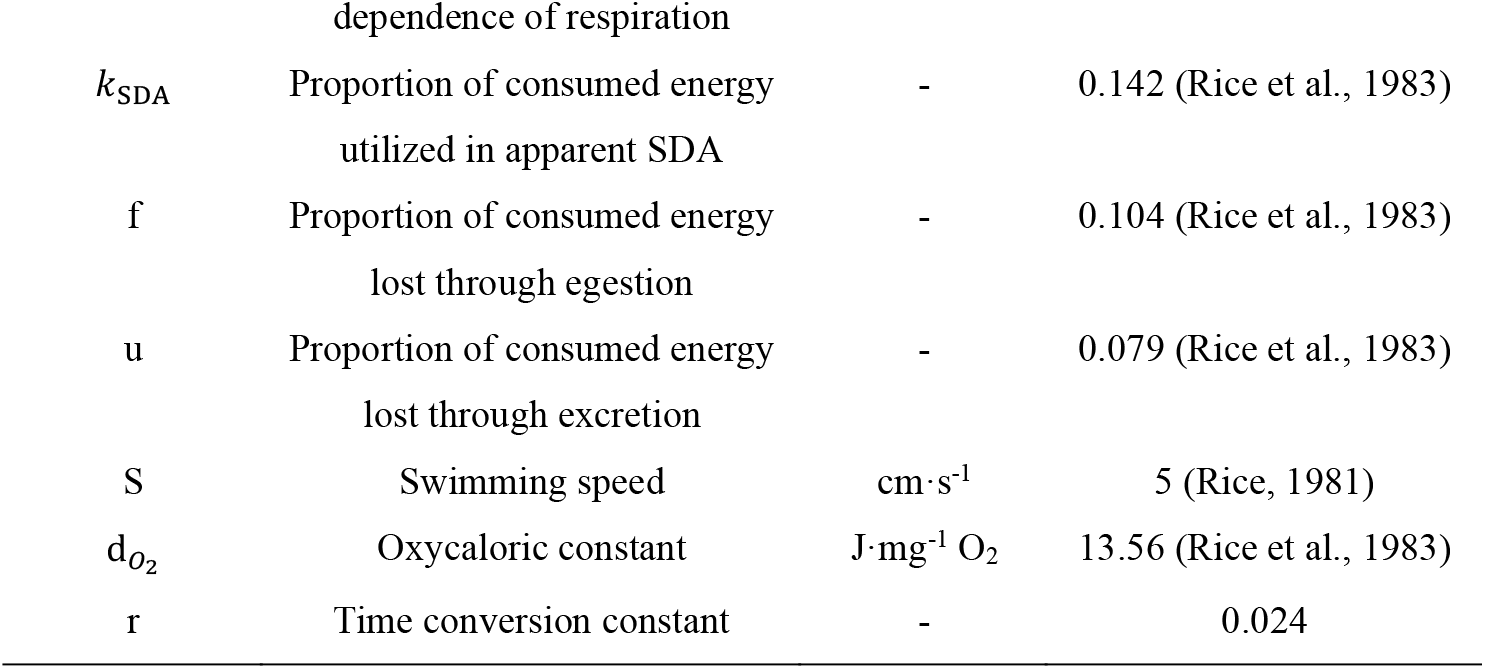
Parameters used in the bioenergetics model.

**Table 3.**
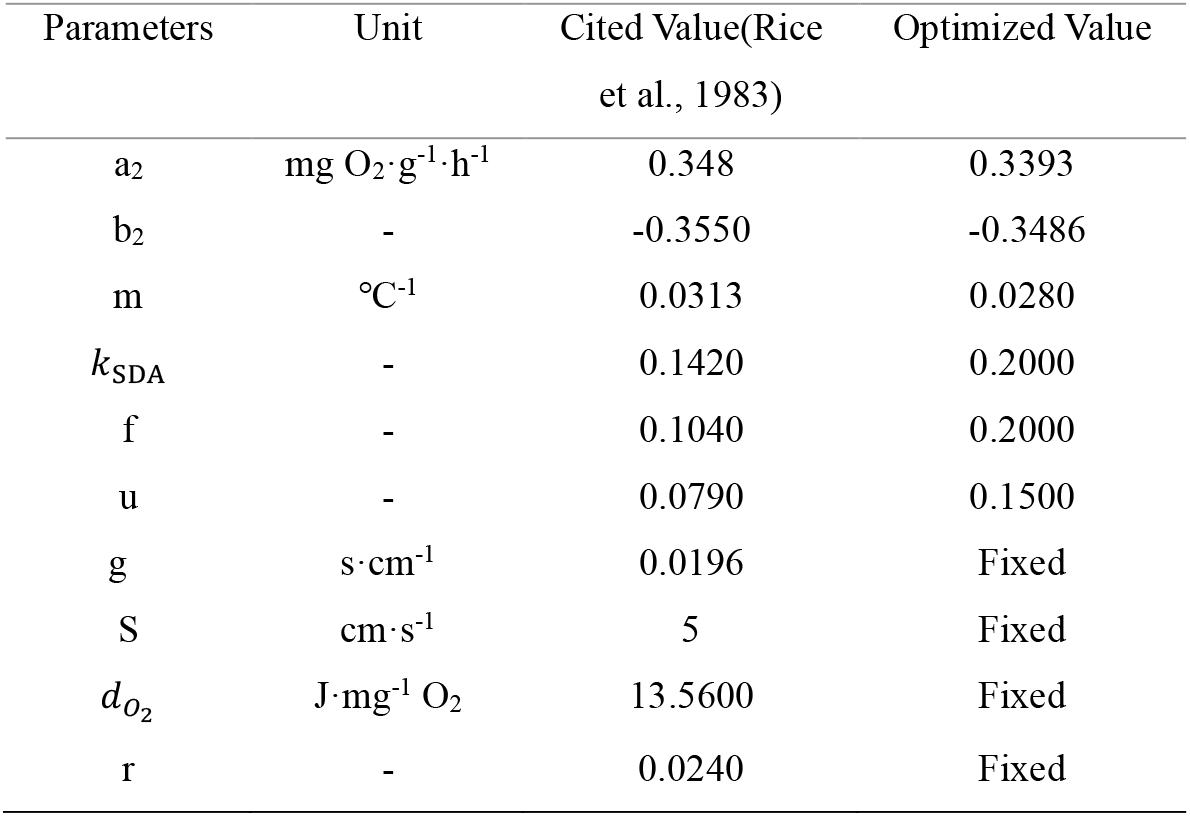
Parameters values from literature and optimization.

**Table 4.**
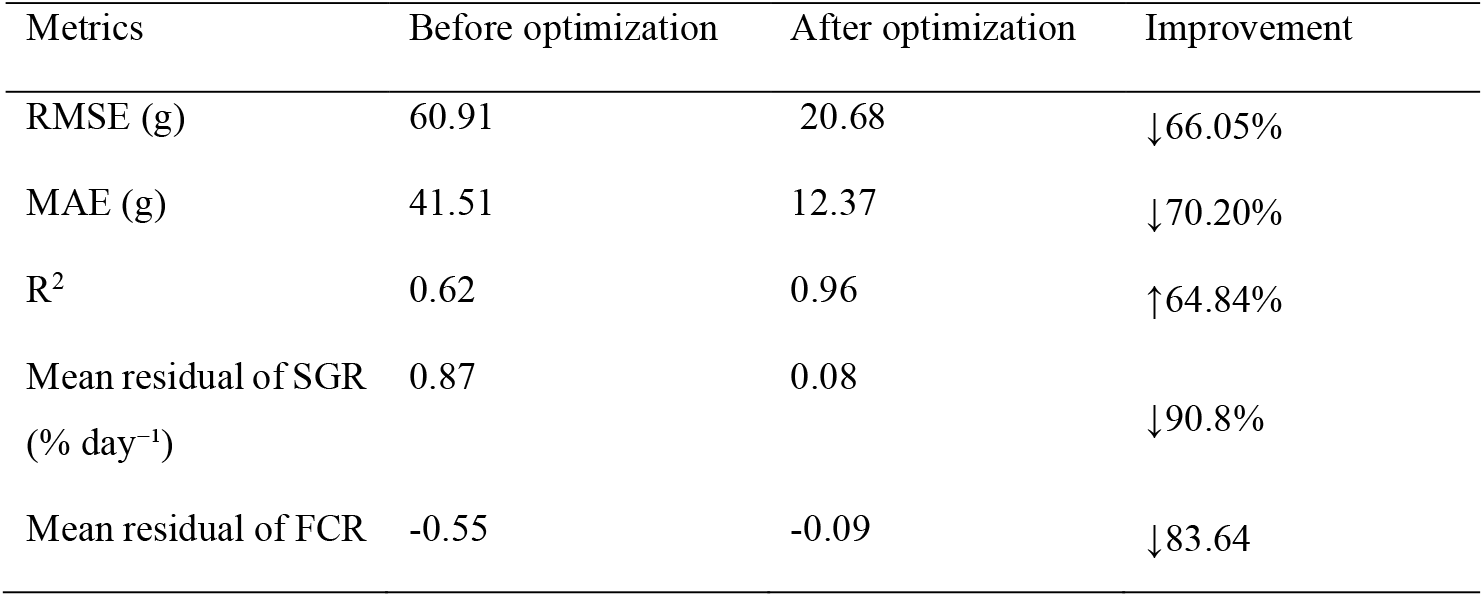
Performance of Bioenergetics Model Before and After Parameter Optimization (Literature-Based Dataset)

**Table 5.**
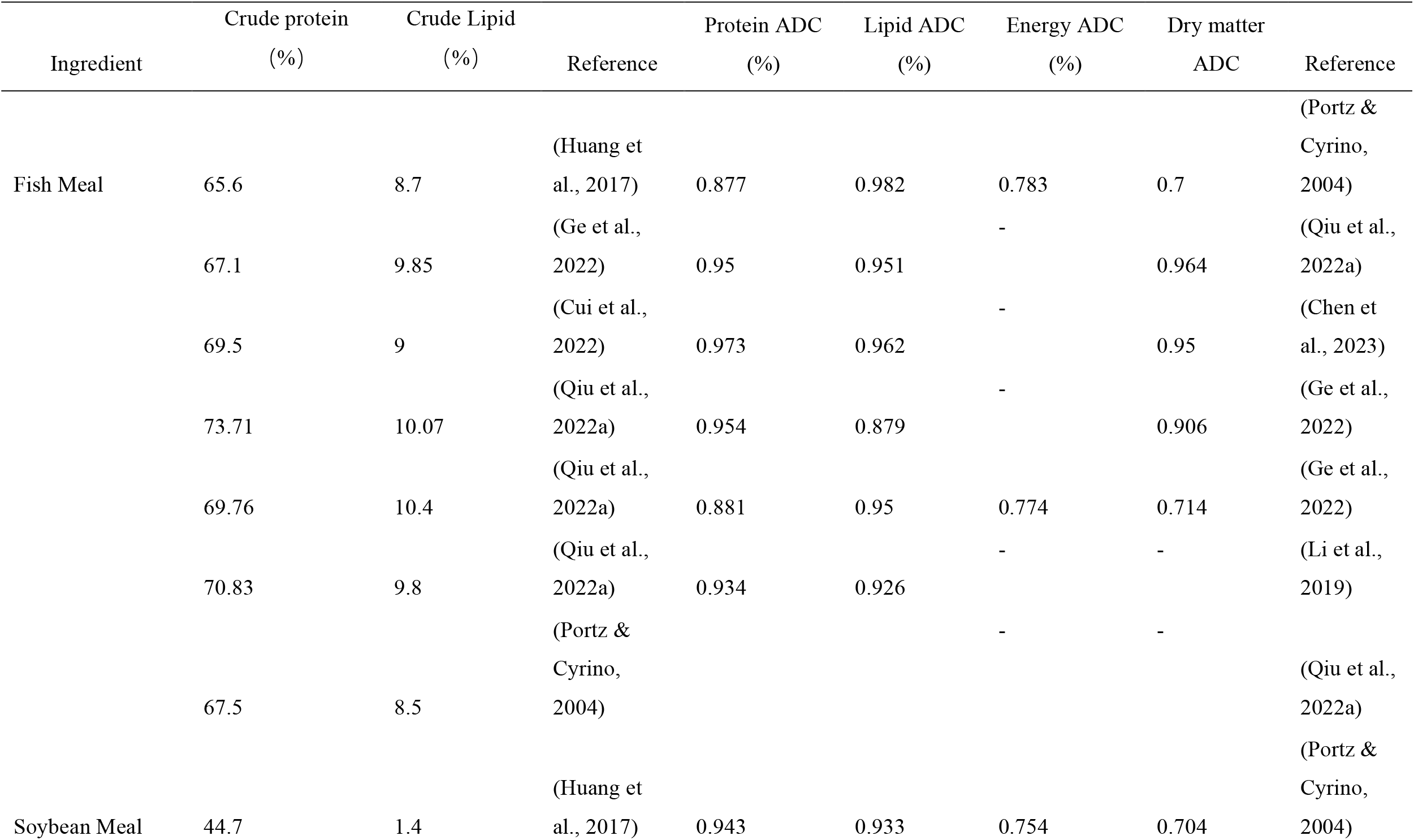
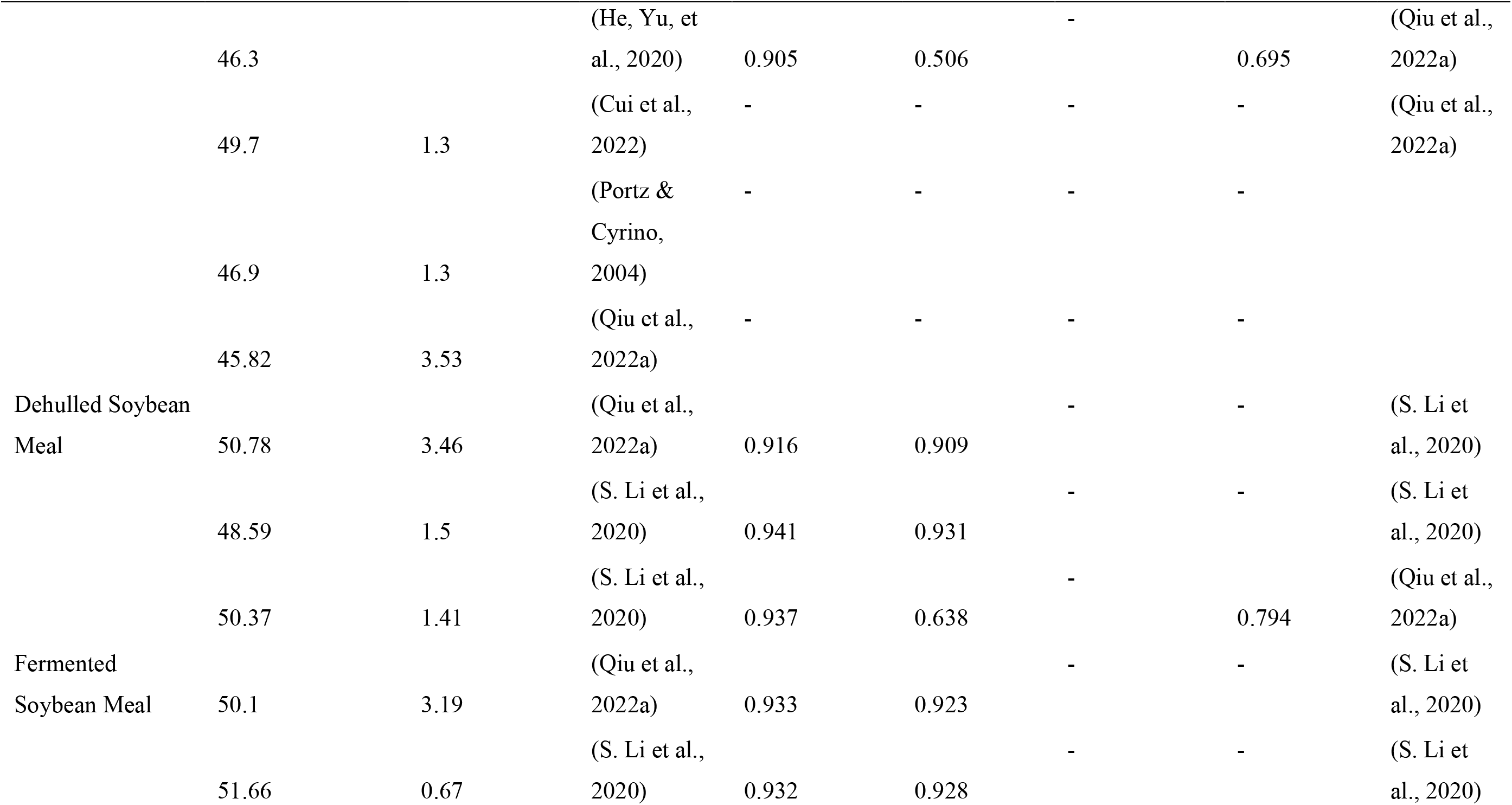
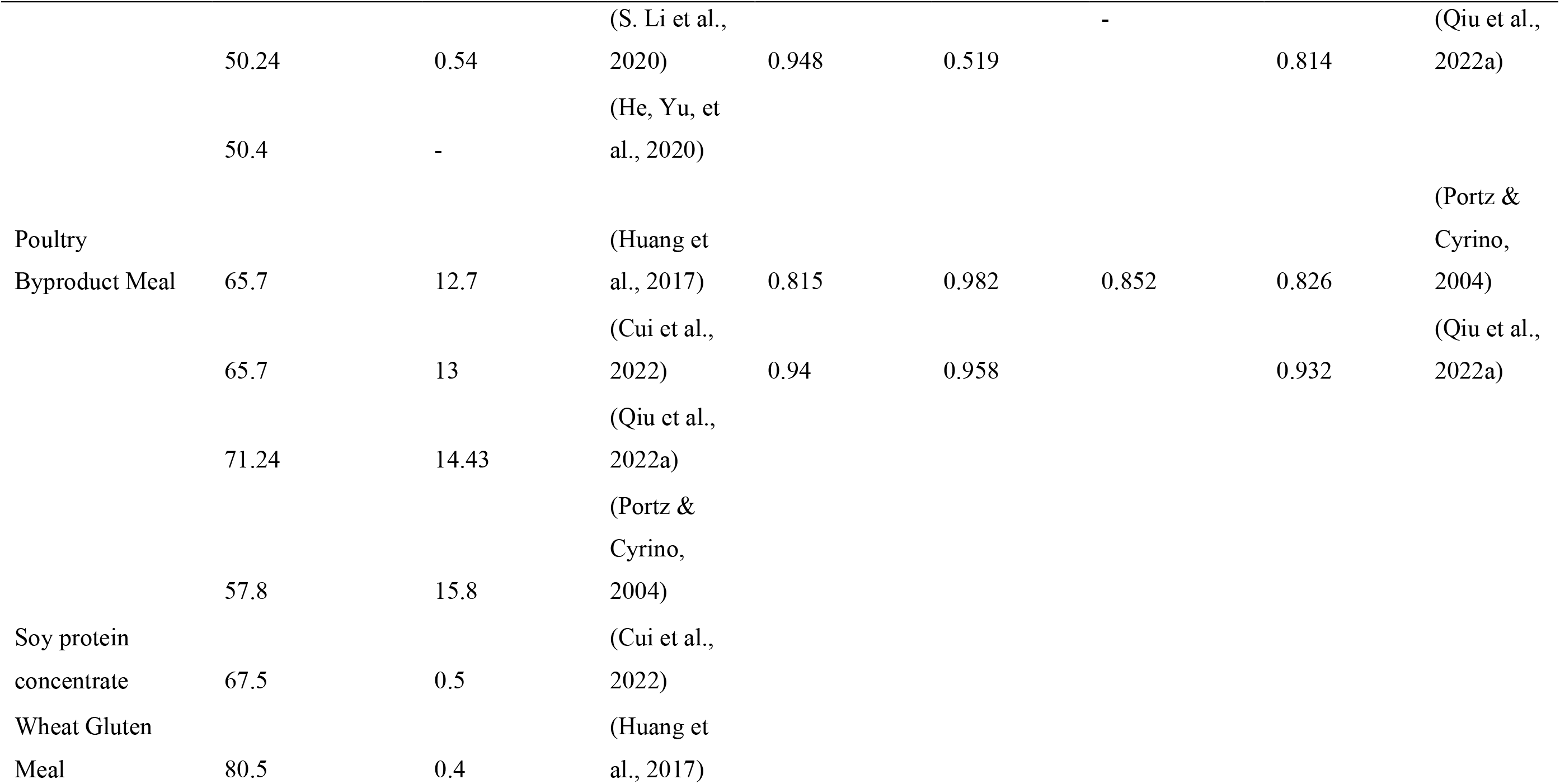
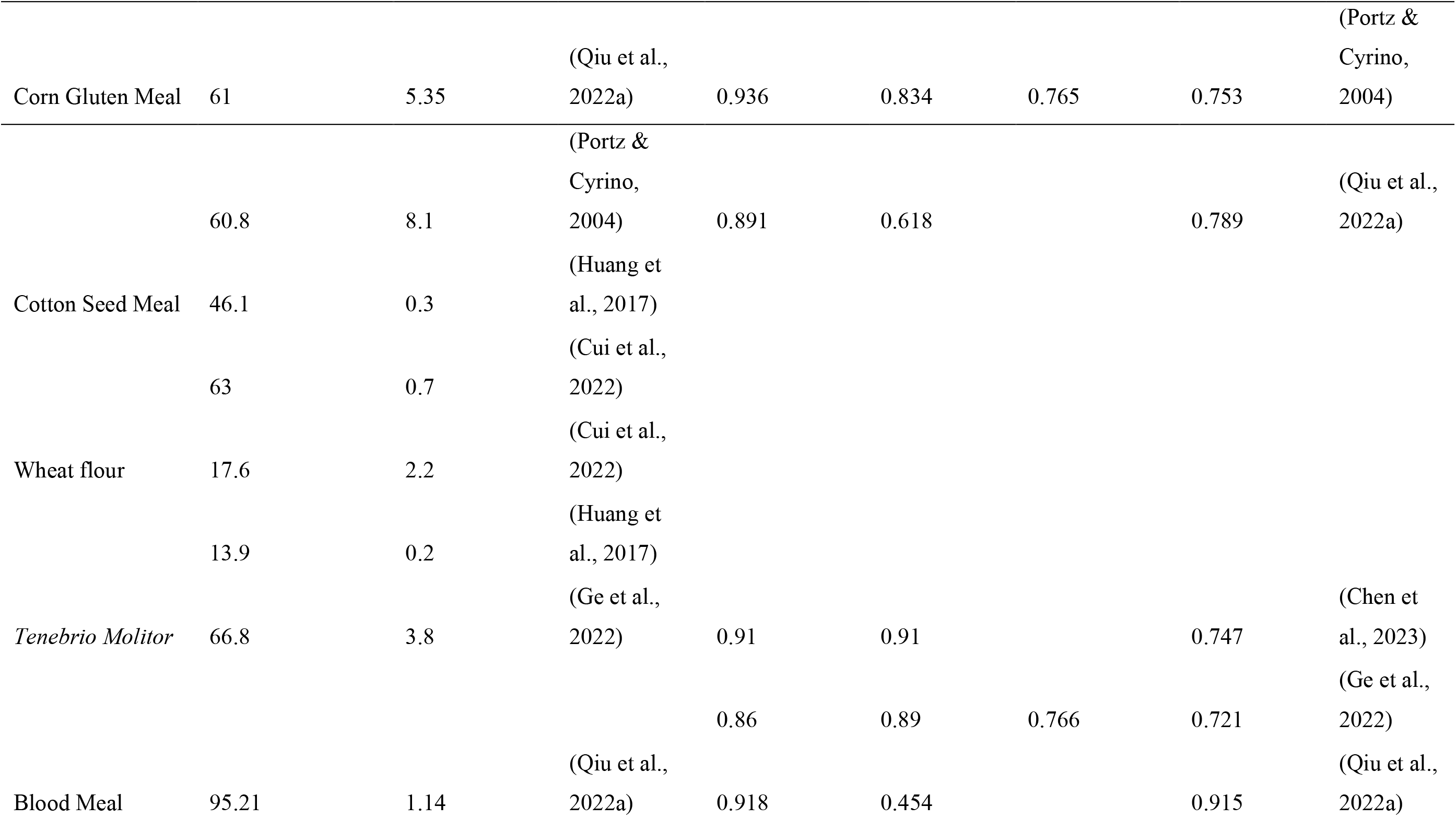
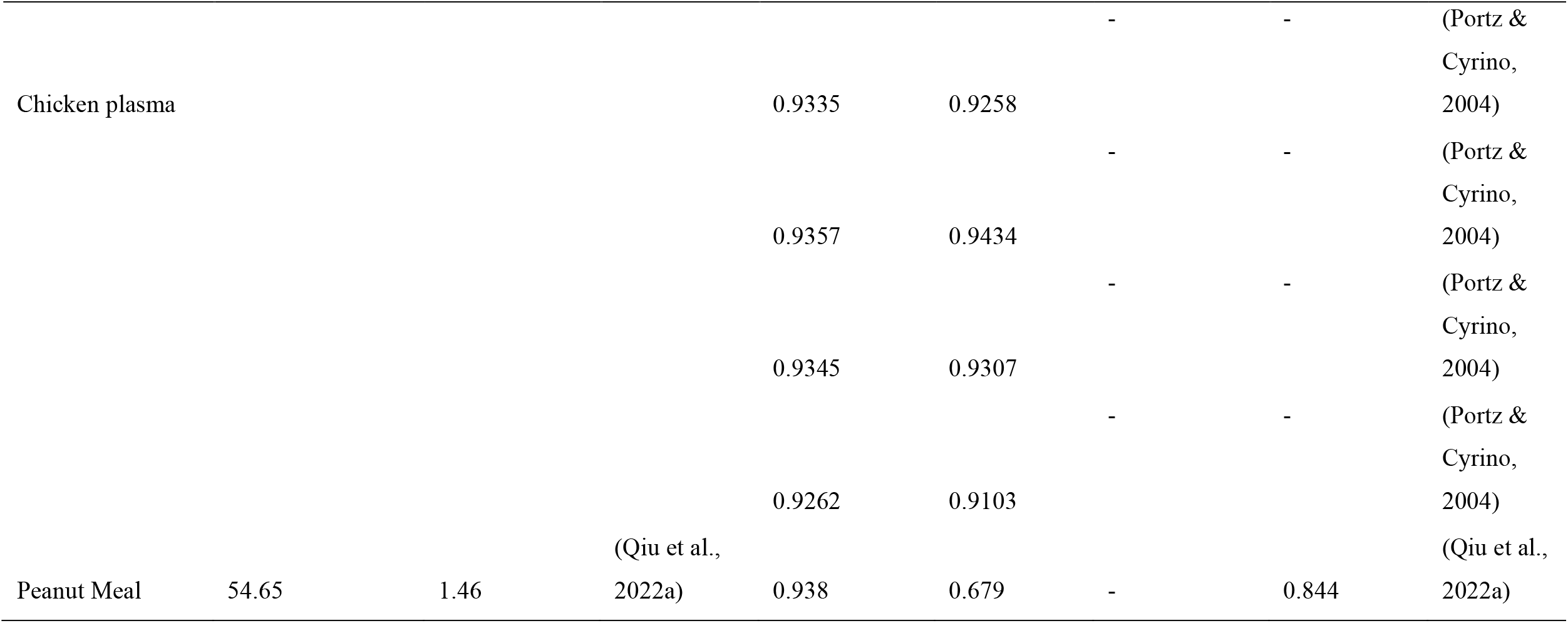
Apparent digestibility coefficients (ADCs) for representative feed components used in energy intake estimation.

**Table 6.**
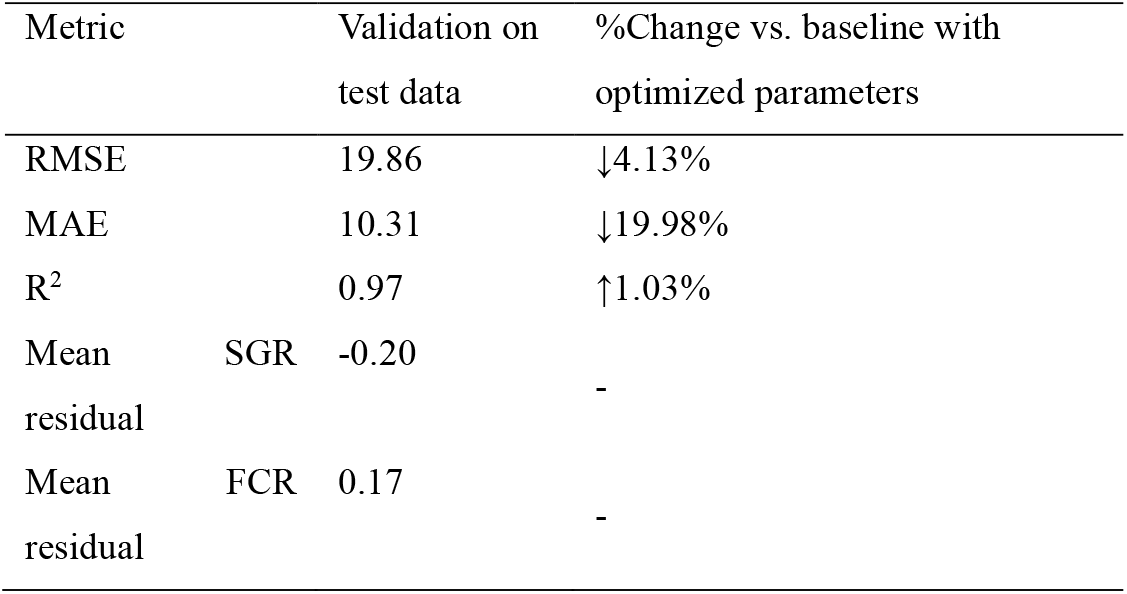
Model outcomes of refined bioenergetics model on literature-based dataset.

**Table 7.**
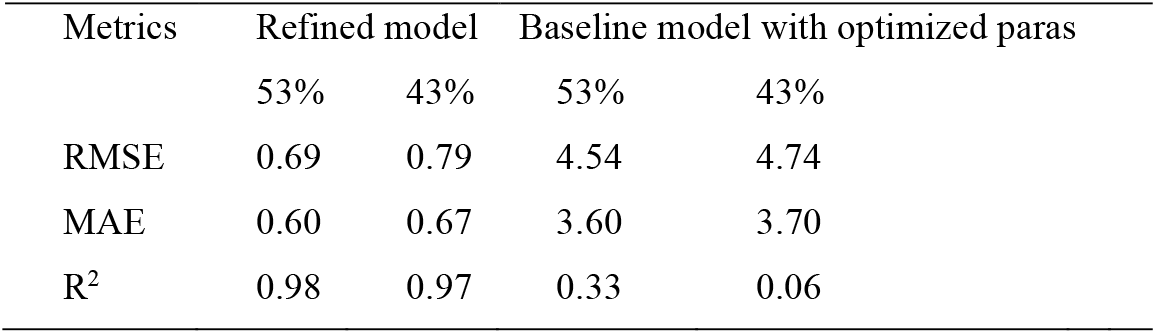
Experimental validation.

## ii. Appendix: Figures

**Fig. 1.**
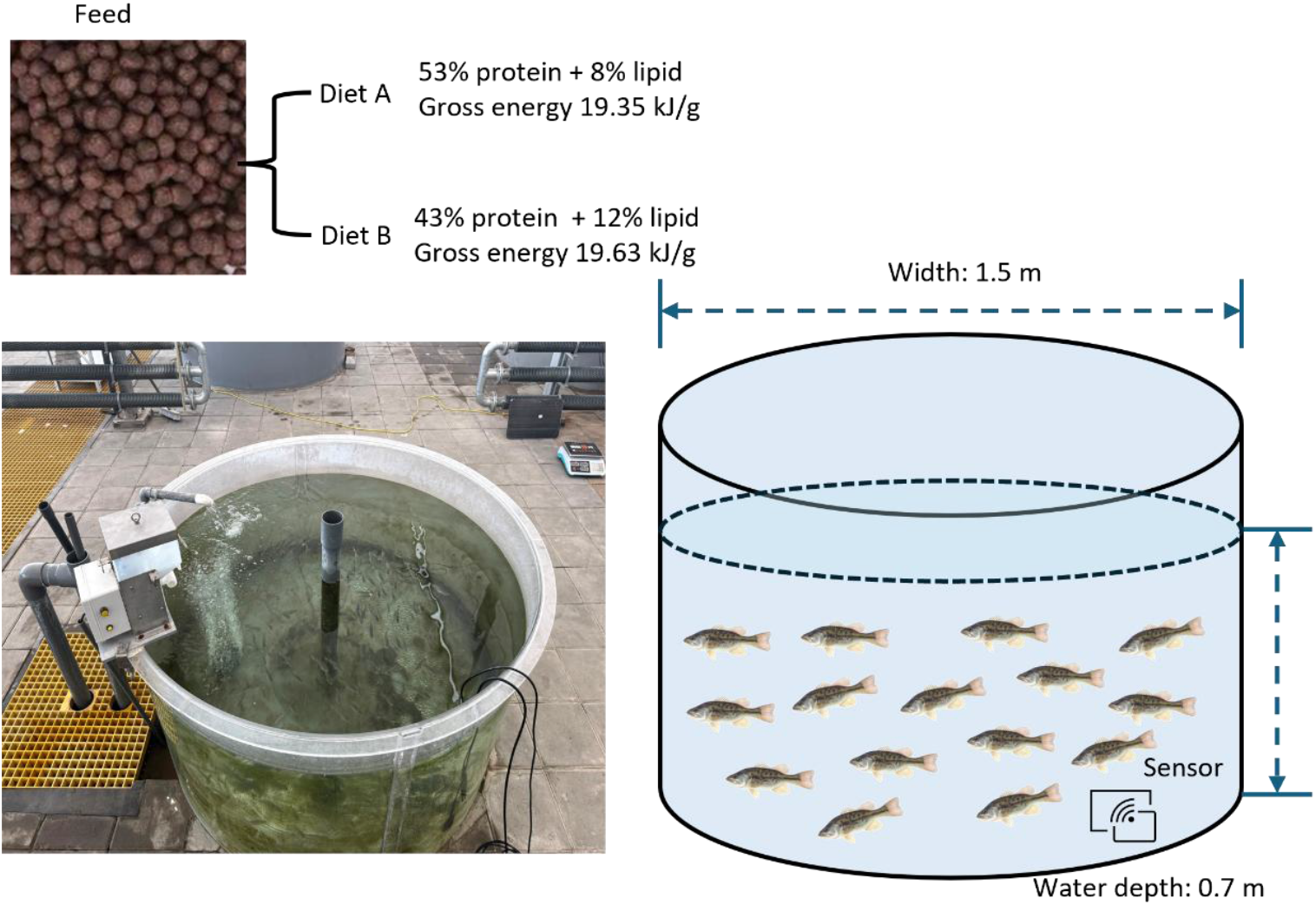
Field experimental design of the largemouth bass RAS trial with two isoenergetic (gross energy) diets

**Fig. 2.**
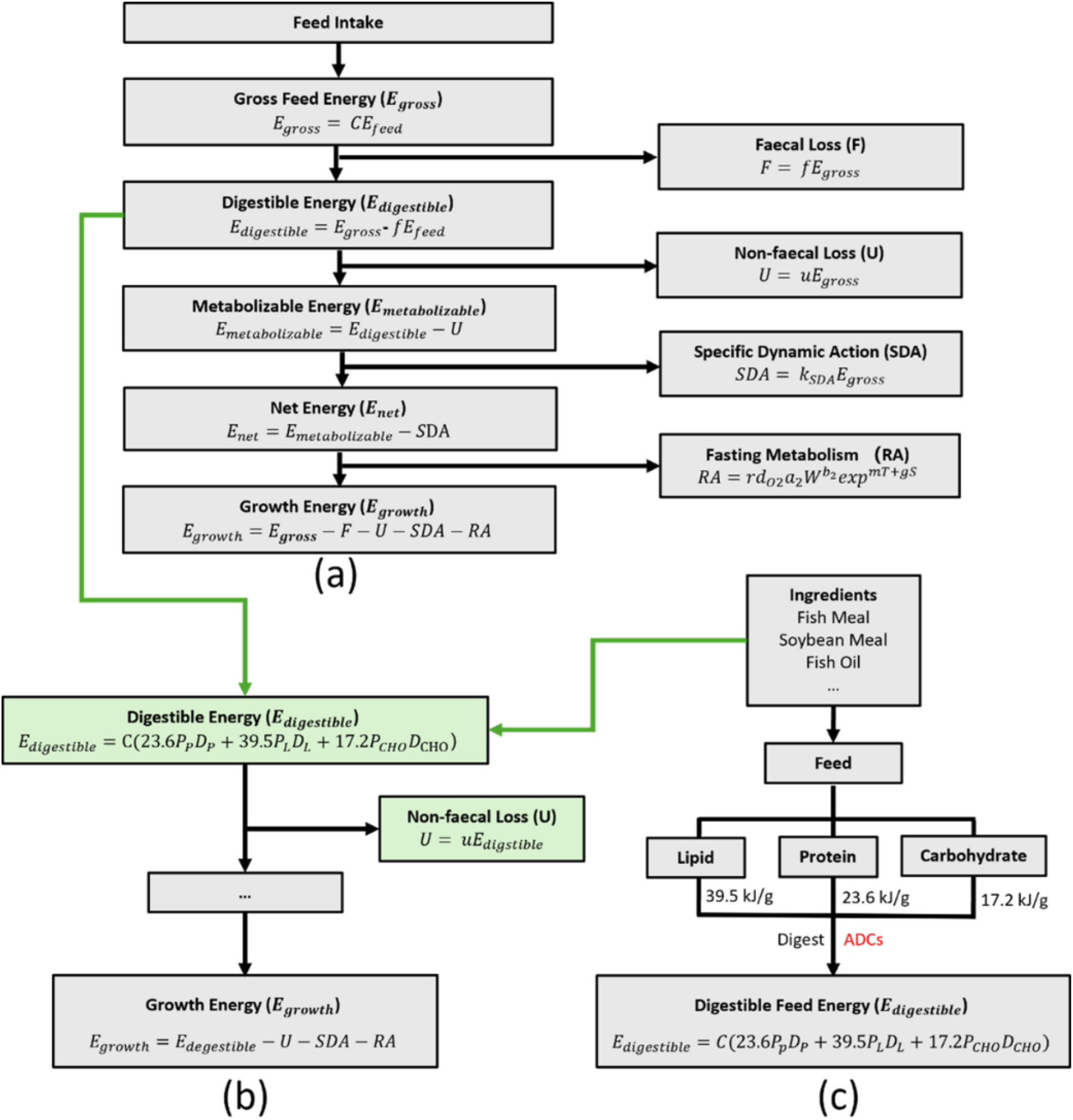
Energy partitioning structure of the baseline (a) and nutrient-explicit refined (b) bioenergetics models, and calculation framework of digestible feed energy (c) based on nutrient composition and ADCs.

**Fig. 3.**
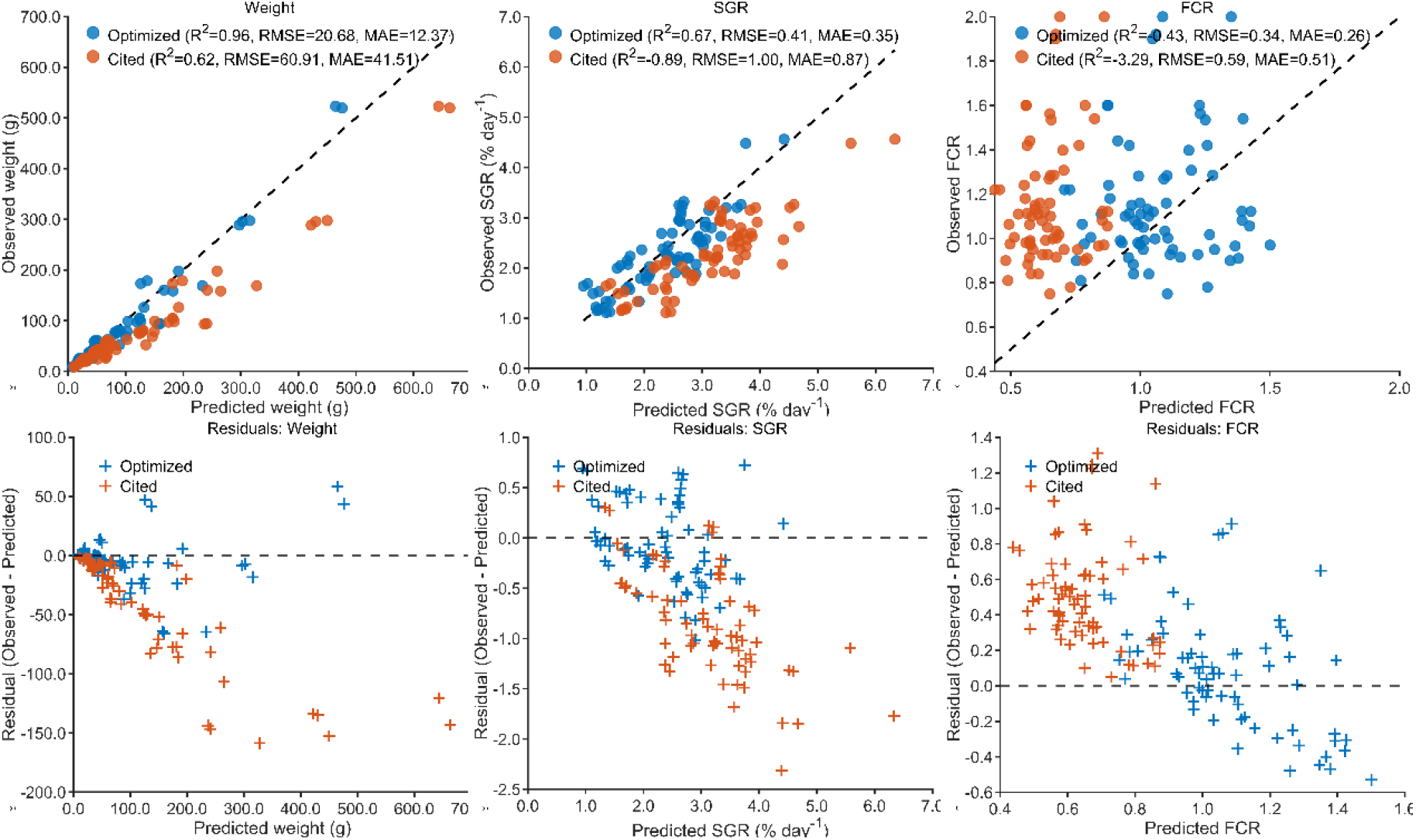
Evaluation of the baseline bioenergetics model: predicted vs. observed values(a-c) and residuals (d-f) for body weight, SGR, and FCR (n=70).

**Fig. 4.**
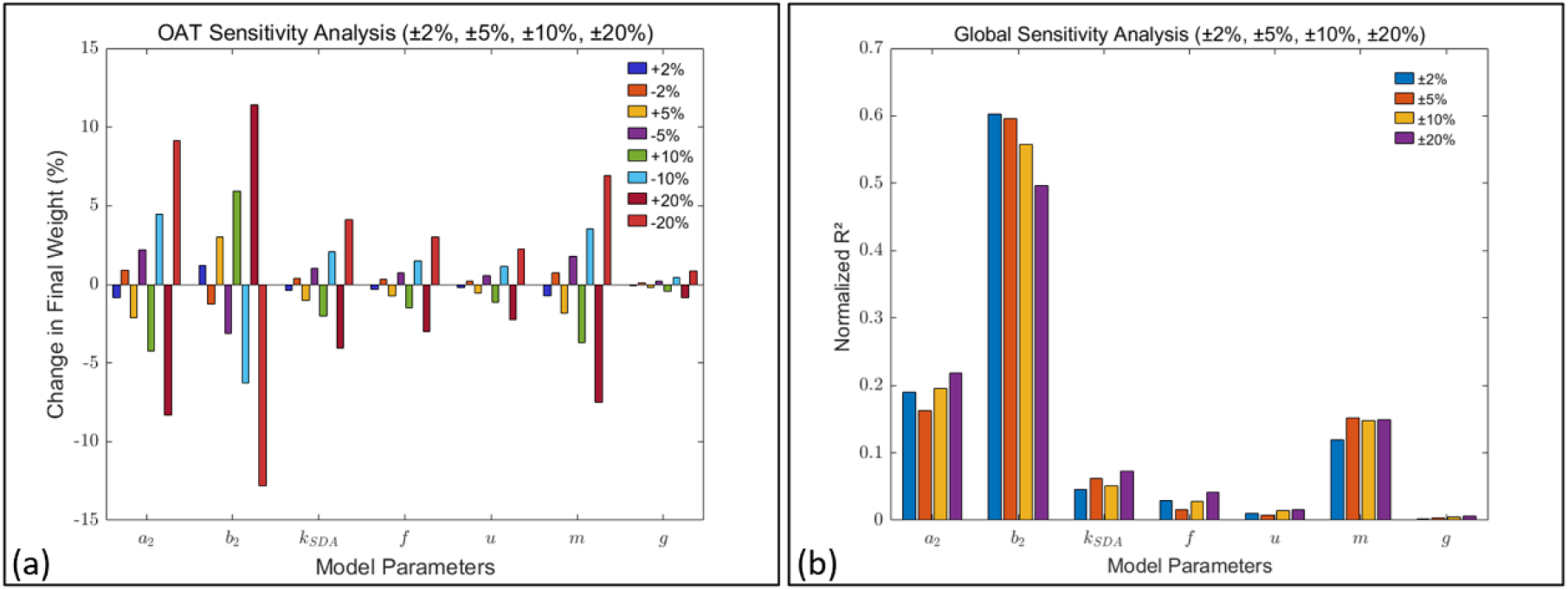
Sensitivity analysis of baseline bioenergetics model by (a) OAT and (b) Monte Carlo Sampling and Normalised Pearson Correlation Coefficients

**Fig. 5.**
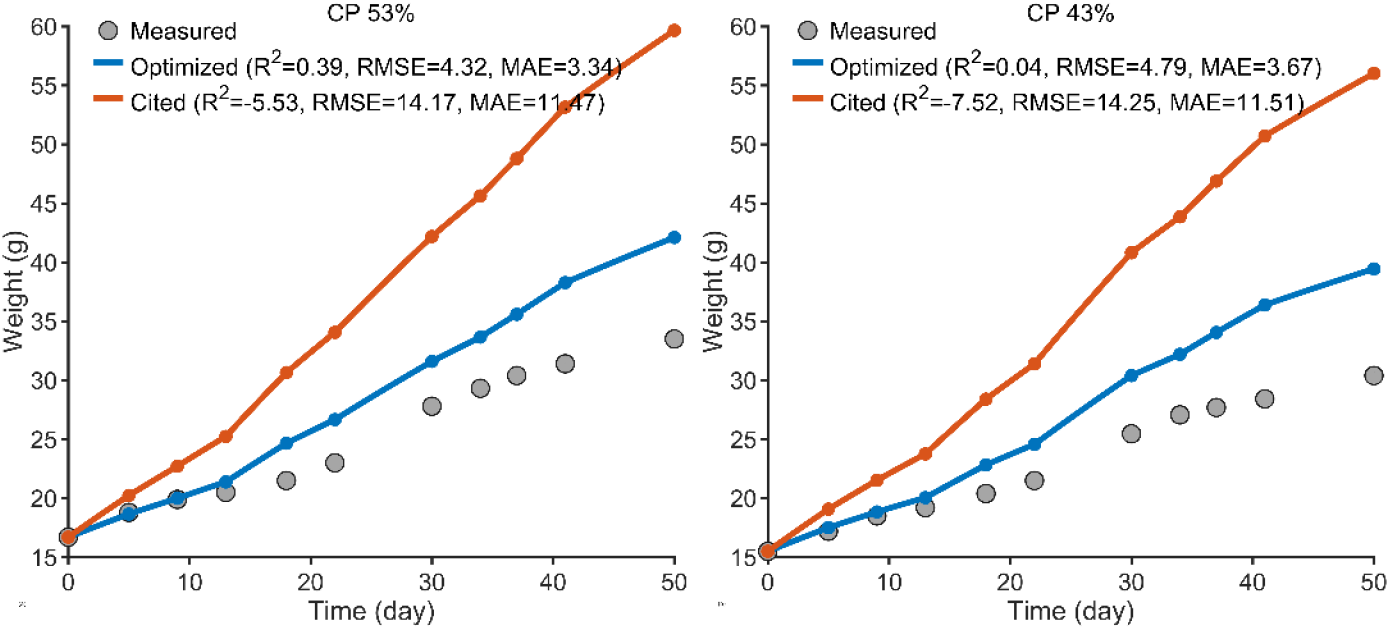
Comparison of baseline model simulations on field data (Diet A and Diet B) before (a–b) and after (c–d) optimization

**Fig. 6.**
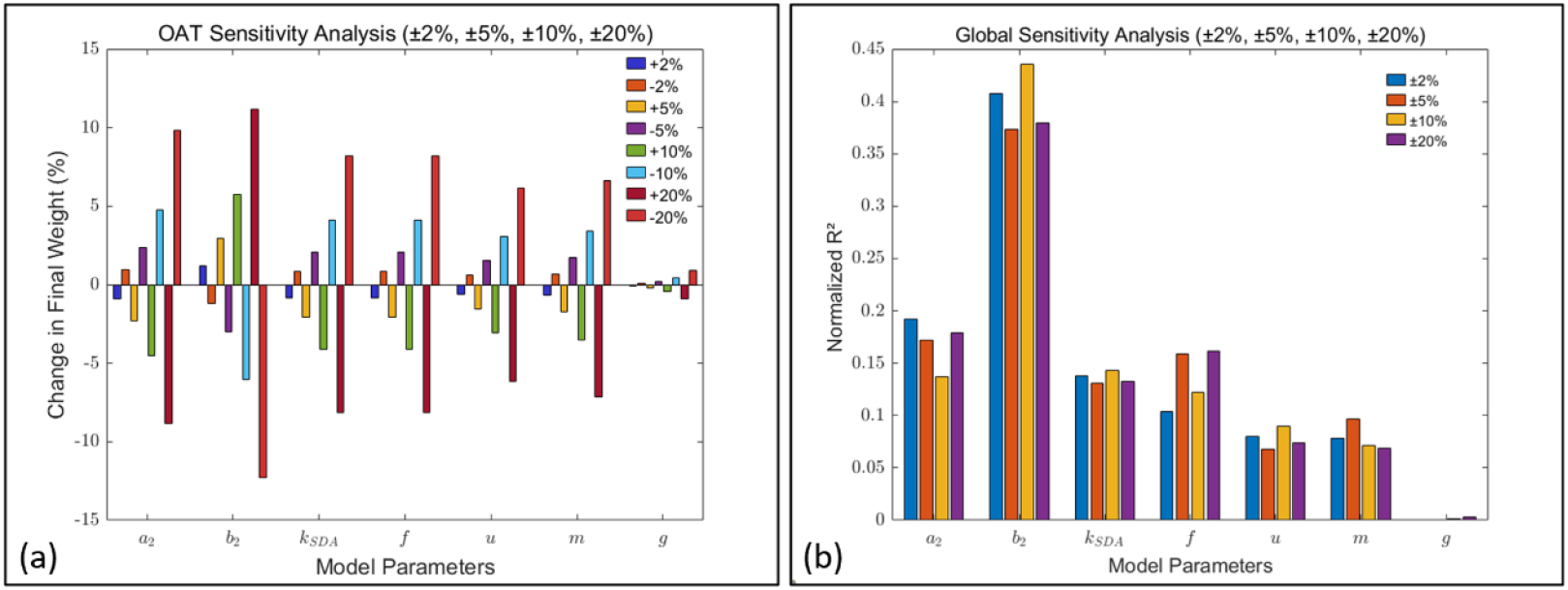
Sensitivity analysis of baseline bioenergetics model with optimized parameters by (a) OAT and (b) Monte Carlo Sampling and Normalised Pearson Correlation Coefficients

**Fig. 7.**
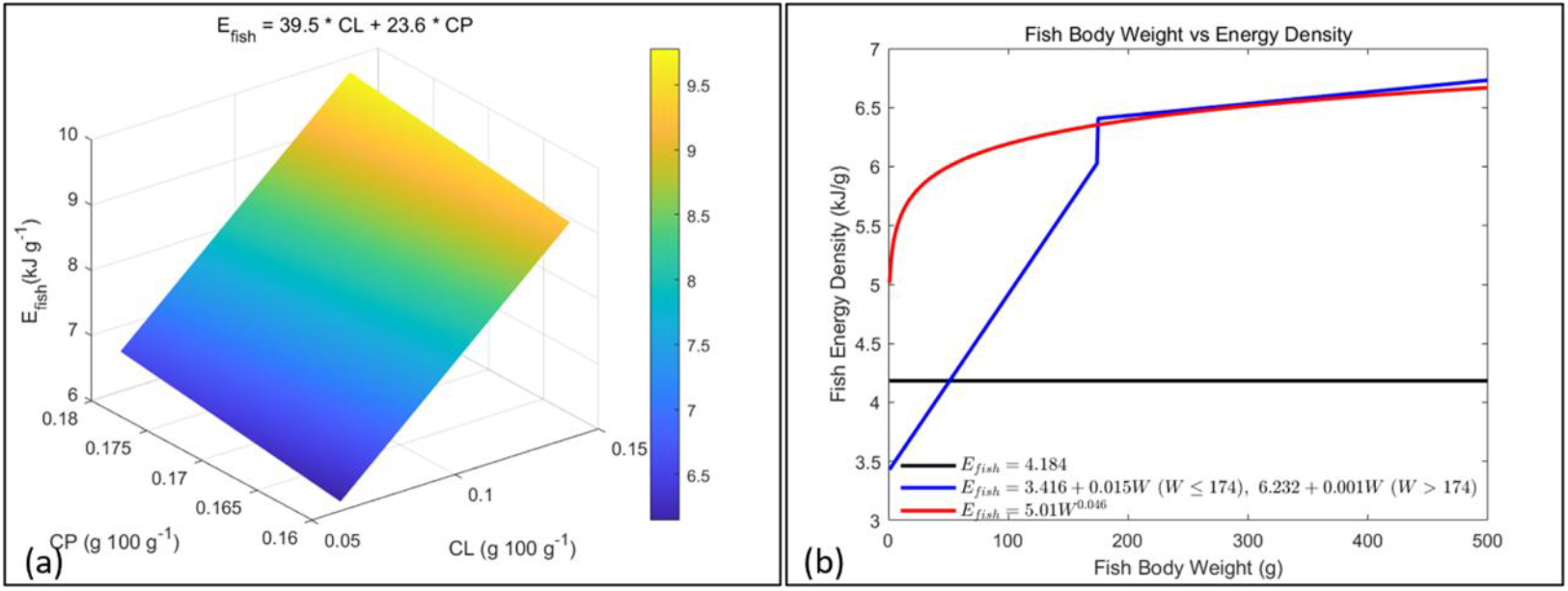
Different methods for adjusting fish energy density

**Fig. 8.**
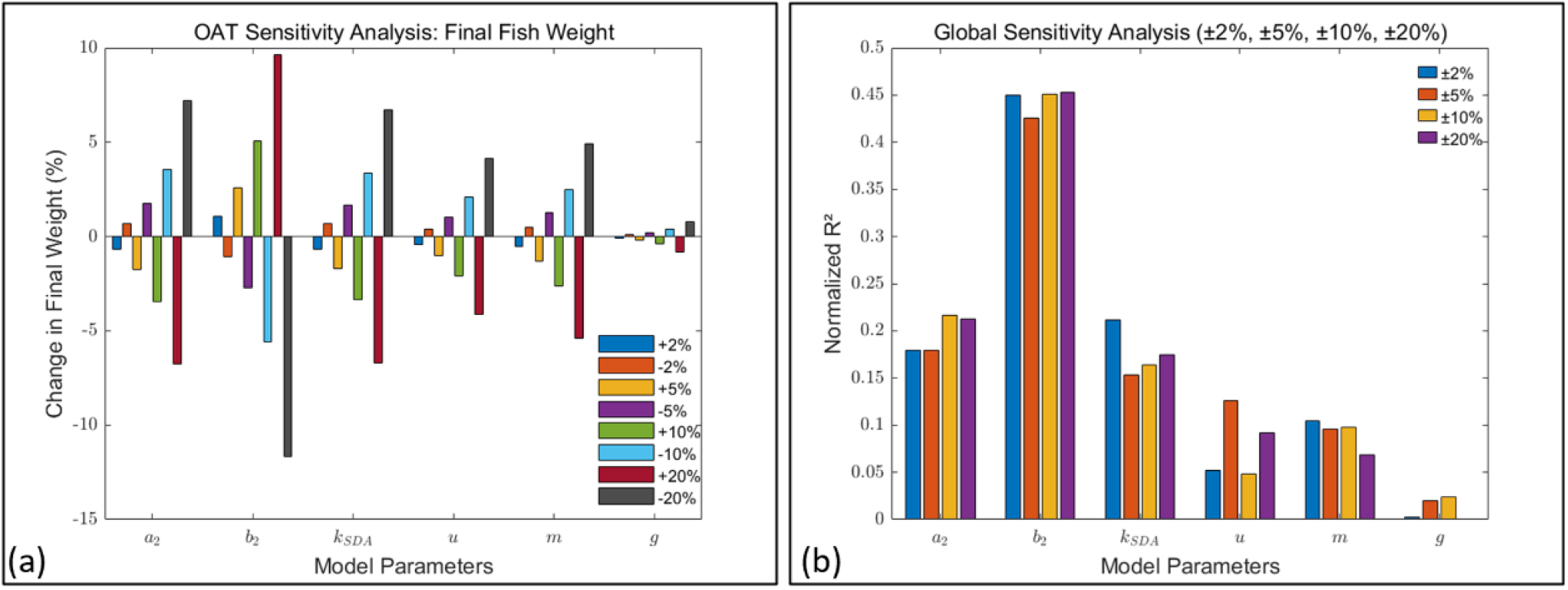
Parameters sensitivity analysis of refined bioenergetics model by (a) OAT and (b) Monte Carlo Sampling and Normalized Pearson Correlation Coefficients

**Fig. 9.**
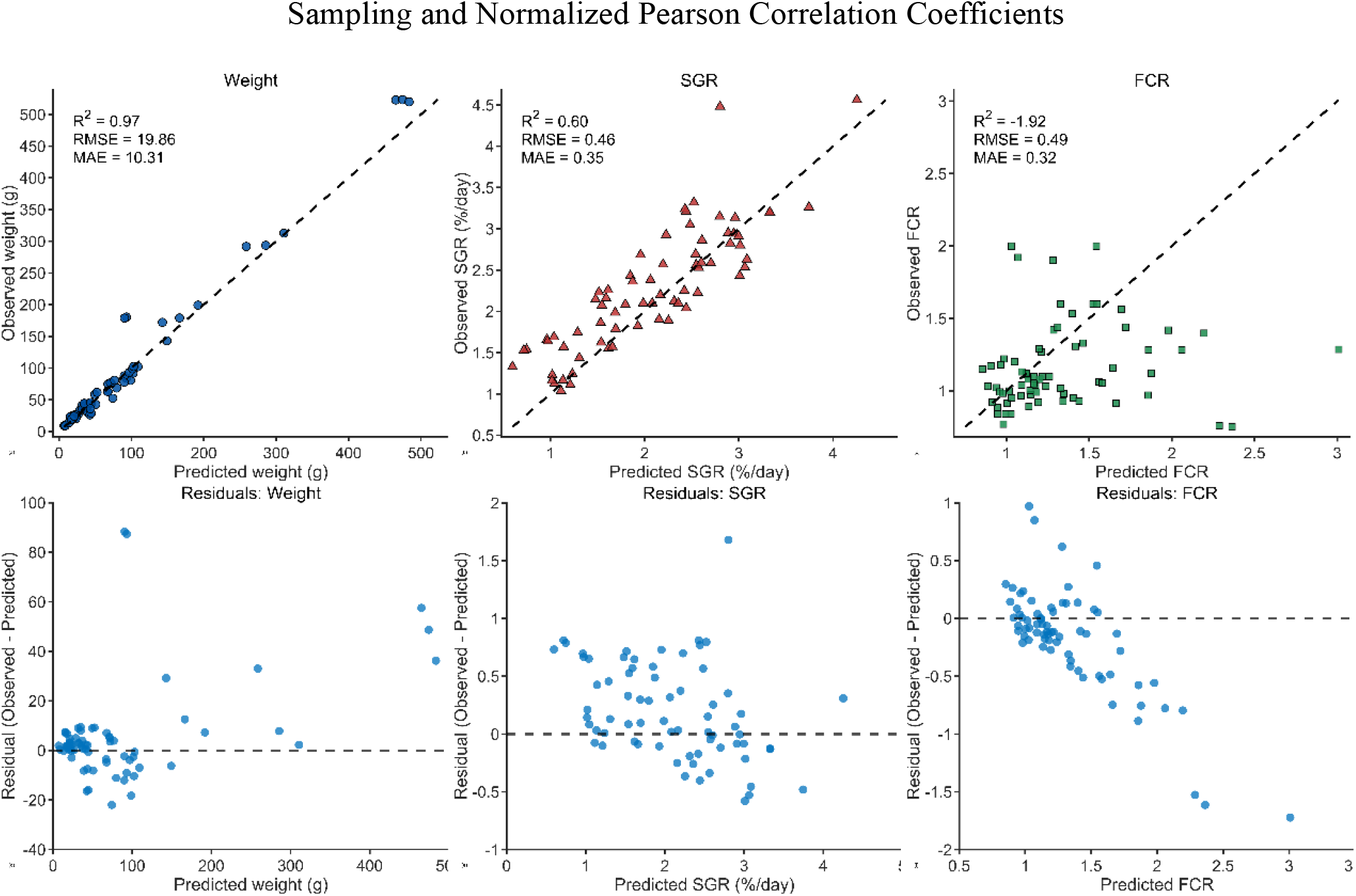
Evaluation of the refined bioenergetics model: predicted vs. observed values(a-c) and residuals(d-f) for body weight, SGR, and FCR (n=70).

**Fig. 10.**
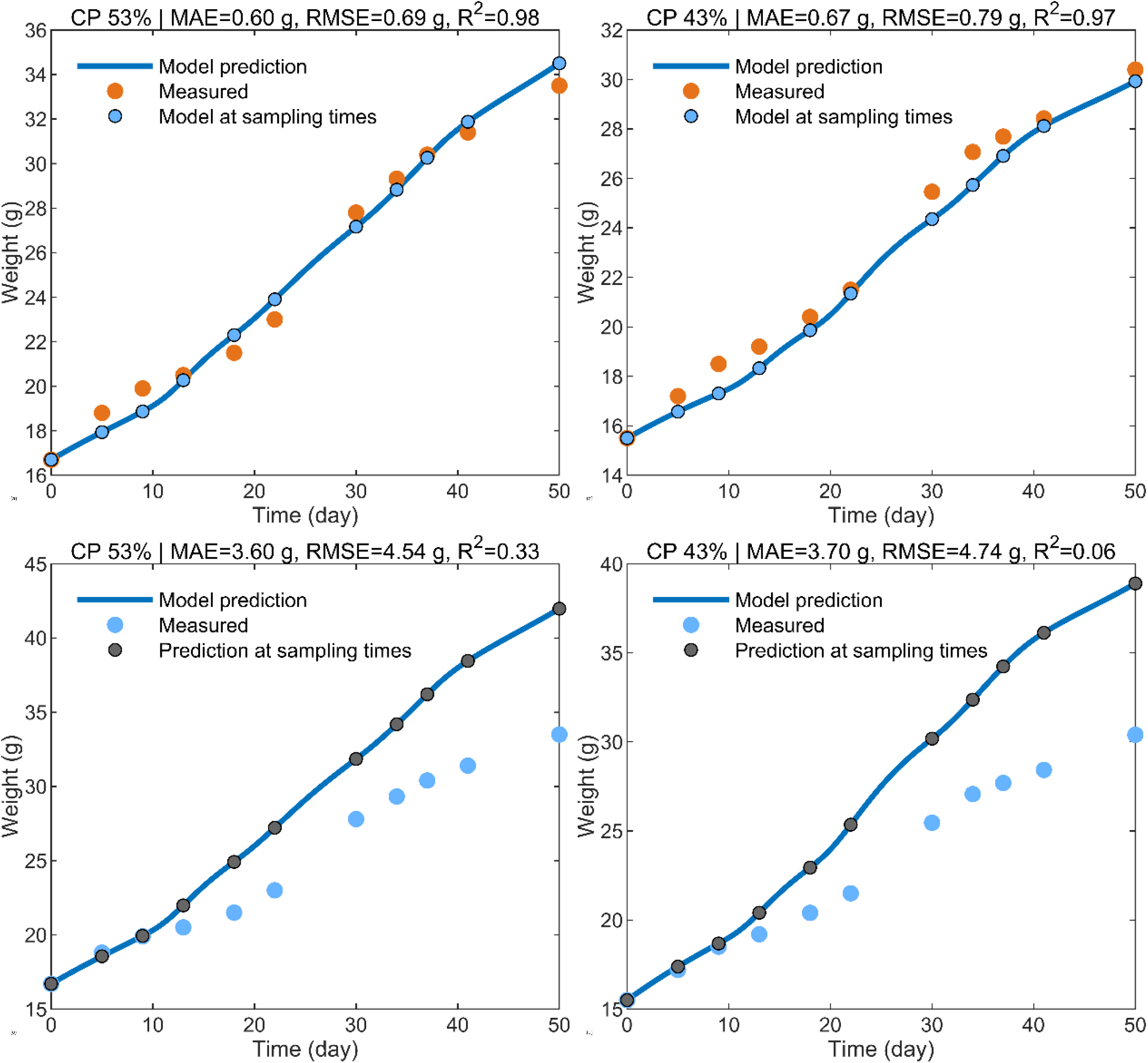
Predicted vs. observed body weight of largemouth bass under two diets using the refined model (a-b) and the baseline model with optimized parameters (c-d).

